# Nuclear pore complex distribution on the nuclear envelope: insights into curvature, chromatin, and actin contributions

**DOI:** 10.1101/2025.01.09.632019

**Authors:** Hugo Lachuer, Orestis Faklaris, Sylvie Hénon, Fabien Montel, David Pereira

**Affiliations:** Université Paris Cité, CNRS, Institut Jacques Monod, 75013 Paris, France; MRI, Biocampus, University of Montpellier, CNRS, INSERM, Montpellier, France; Laboratoire Matière et Systèmes Complexes, Université Paris Cité, CNRS, UMR7057, 10 rue Alice Domon et Léonie Duquet, F-75013, Paris, France; Laboratoire de Physique, UMR CNRS 5672, ENS de Lyon, Université de Lyon, Lyon, France

## Abstract

The Nuclear Pore Complex (NPC) is an ancestral feature of eukaryotic cells, essential for the exchange of material and information between cytoplasm and nucleoplasm. While, typical eukaryotic cells harbor thousands of NPCs covering the nucleus surface, a quantitative characterization of their spatial distribution remains lacking. Taking advantage of super-resolution microscopy, we imaged and segmented individual NPCs in myoblast cells to reconstruct the nuclear surface. Our spatial statistics analysis of NPCs on the nuclear surface reveals the existence of a 400nm repulsion length between NPCs, likely reflecting the mesh size of the underlying lamin network. Furthermore, we identify three key correlates of NPC density. (1) Myoblast actin fibres, which can indent the nucleus, correlate with a decrease in NPC surface density, contrary to what was reported in previous studies; (2) Regions of the nuclear envelope near large heterochromatin domains exhibit reduced NPC surface density; and (3) There is a non-monotonous relationship between NPC density and the nuclear envelope curvature. This last observation is consistent with the existence of NPC-preferred curvature suggesting curvature sensing properties. In conclusion, this study provides new insights into the spatial distribution of NPCs, shedding light on possible new determinants of NPC spatial distribution.

## Introduction

The only gateway between the nucleus and the cytoplasm of eukaryotic cells is the Nuclear Pore Complex (NPC). It is a complex formed of multiple copies of more than 30 different proteins known as nucleoporins, forming an octameric structure^1^. Typical eukaryotic cells contain thousands of NPCs, with their number increasing as the cell cycle progresses^2^. NPCs are assembled though two main mechanisms: by post-mitotic assembly, where they bind to chromatin and integrate into membrane fenestrae to form the Nuclear Envelope (NE) at the end of mitosis, and by an “inside out” mechanism during interphase, in which the NPC complex nucleation occurs on the nucleoplasm side of the NE, leading to the fusion of inner and outer nuclear membranes^3^. Despite their critical functions, the spatial organization of NPCs remains poorly known.

Once assembled, NPCs are mobile in yeast models as demonstrated by yeast karyogamy experiments where a nucleus with fluorescent NPCs fuses with an unlabeled nucleus^4^. However, in mammal models, NPCs are anchored to the nuclear lamin and chromatin, hindering their mobility^5^. However, it has been recently show that Gaussian curvature of NE could dilute lamins^6^, potentially weakening the NPC anchoring and increasing their mobility. Despite their low mobility, particular spatial organizations of NPCs have been already reported in various studies^7–10^. Yet, despite the growing body of super-resolution data, little is known about the spatial organization of NPCs and how the surrounding environment influences it.

An important aspect of NPC organization is how it responds to physical constraints on the nucleus, which arise during physiological processes such as cell migration or muscle contraction^11^. These mechanical deformations of the nucleus are known to affect its physiology, for example, physical cues have been shown to induce the translocation into the nucleus of transcription factor co-activators YAP/TAZ and MRTFA, and of the lysine methyltransferase SMYD3^12–14^. Many of these physical constraints on the nucleus are generated by the cytoskeleton. Actin cables, for example, can be mechanically linked to the nucleus by Nespring-2G and SUN2 forming the LINC (Linker of Nucleoskeleton and Cytoskeleton) complex. Actin bundles can also associate with the nucleus through a series of LINC complexes to form a Transmembrane Actin-associated Nuclear (TAN) line, which leads to nuclear indendation^15^. This indentation has been shown shown to trigger nuclear polarization^16^ and is thought to be associated with NPC enrichment^17,18^. This claim, however, is primary based on co-localization measured by classical confocal microscopy, which may overlook local geometrical effects. Specifically, TAN-line indentation generates a curvature of the NE, increasing the membrane area assessed and potentially leading to the misinterpretation of signal enrichment when projected on the x-y plane. This caveat underscores the importance of using super-resolution microscopy to accurately account for NE geometry in these indented regions.

Another key factor that could regulate NPC spatial organisation is the local chromatin state. Given that one of the primary functions of NPCs is to serve as gateways for transcription factors and mRNA, we hypothesized that NPCs might be more densely clustered near active chromatin regions for optimization purposes. While the interplay between chromatin compaction and NPCs^19^ is well-established, the specific spatial relationship between has never yet been investigated.

One last promising candidate to spatially organize NPCs is NE curvature, especially since the NPCs themself exhibit curvature-sensing features. Specifically, an amphipathic α-helical motif (ALPS) responsible for sensing membrane curvature and originally discovered in the Golgi-associated protein ArfGAP1 has been localized in Nup133^20^. Nup133 is part of the Nup107-160 complex located in the equatorial region of the NPC, where the inner and outer nuclear membranes merge. It has been demonstrated that the N-terminal domain of Nup133 binds to liposomes depending on their radius^20^. Moreover, it has been shown, *in vitro*, that both Nup1 and Nup60 transform spherical synthetic liposomes into highly curved membrane structures^21^ and that in yeast, high expression of the amphipathic helices of Nup1/Nup60 causes deformation of the NE^21^. Furthermore, the NPC scaffold may be evolutionary related and architecturally similar to vesicle coats^22,23^, as hNup107 subcomplexes show structural parallels to COPI, COPII, and clathrin proteins^24^. These curvature-sensing features of NPC components may play a key role in the potential accumulation of NPCs in regions of the nucleus with favorable curvature.

In this study, we test the hypothesis that the NPC is sensitive to the aforementioned factors and especially able of sensing the local curvature of the NE. To this end, we examined the spatial organization of NPCs using super-resolution microscopy with advanced numerical methods for surface reconstruction, curvature computation, and spatial statistics. We segmented the 3D positions of NPCs on the nuclear envelope of a murine myoblast cell line (C2C12) from Structured Illumination Microscopy (SIM) images. We reconstructed the nuclear surface from these NPCs and computed the associated curvature as well as the local NPC surface density. Our analysis quantitatively demonstrates that NPCs are not randomly distributed; instead, they are enriched in curved regions and less dense in actin-indented and heterochromatin-dense areas. These findings suggest that the spatial distribution of NPCs is influenced by a complex interplay of multiple factors.

## Results

### Nuclear surfaces can be reconstructed from super-resolved NPC imaging

To investigate the spatial distribution of NPCs on the NE, we reconstructed nuclear surfaces from NPC coordinates. To this end, we seeded C2C12 myoblast murine cell line onto rectangular adhesive micro-patterns. The patterning of C2C12 promotes the formation of TAN lines that indent the nucleus, thus creating physical constraints thought to affect NPC distribution^17,18^. We therefore imaged NPCs in our C2C12 model by super-resolution SIM microscopy (**Fig. 1A**). The lateral resolution of the imaging (Point Spread Function (PSF) Full Width at Half Maximum) was estimated to be 110nm (using sub-resolution fluorescent beads in our experimental conditions) and the axial resolution estimated to be 300-350nm^25,26^.

**Figure 1.**
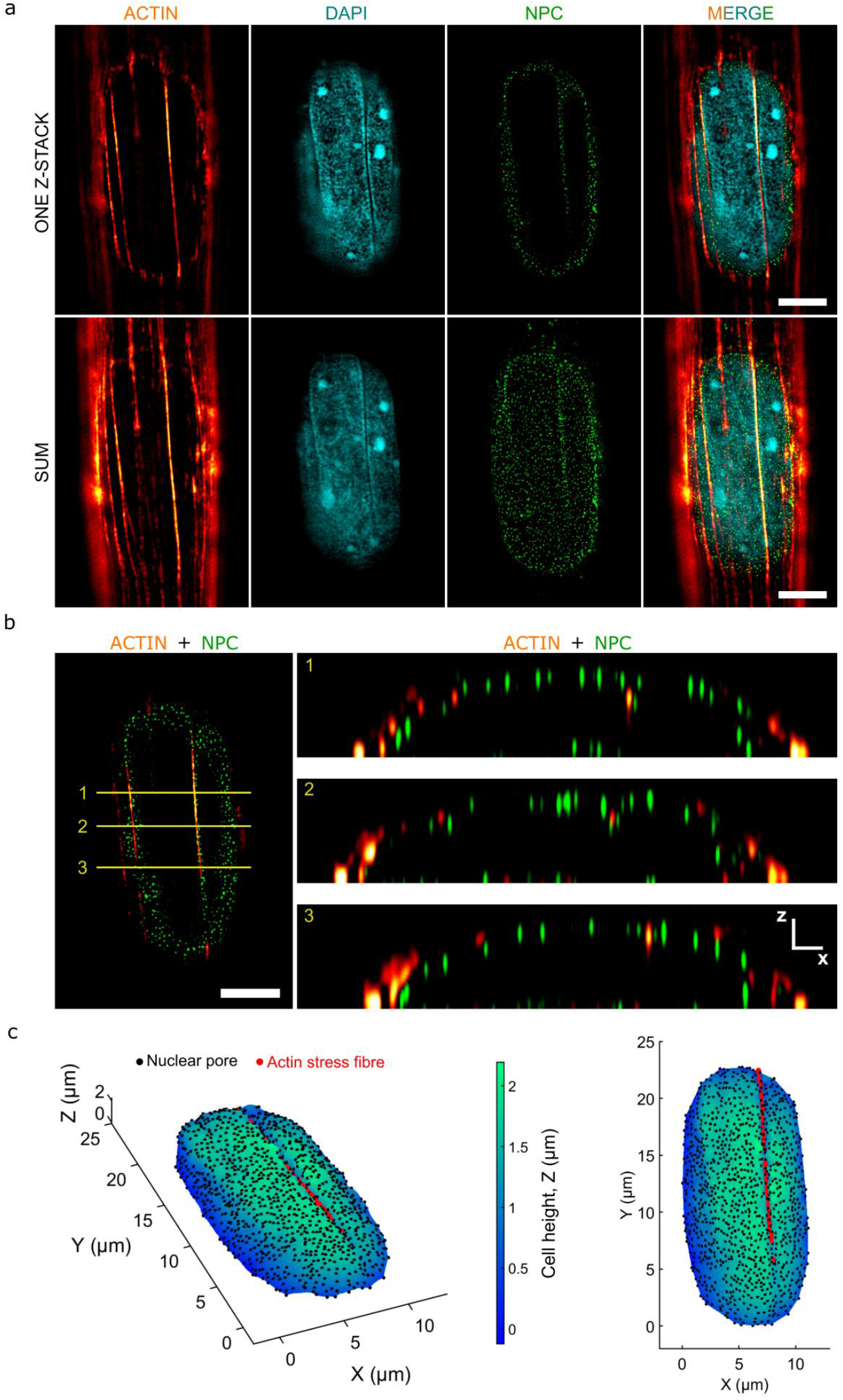
Nuclear surface reconstruction by super-resolution imaging of NPCs. **a**. SIM images of C2C12 seeded on line micropattern. Actin is stained in orange (SiR-actin), NPC in green (WGA staining) and chromatin in blue (DAPI). Single z plane on upper line and z-projection on the lower line. Scale bar is 5µm. **b**. Left image shows 3 lines for x-z cross-sections displayed on the right side. Scale bar is 5µm for the xy image and 1µm (in x and z directions) for the xz images. **c**. Surface reconstruction by interpolation of the NPCs from the nucleus presented in a. Black dots represent segmented NPCs, red dots the actin fibre and color code represents the nucleus height z.

Clear nuclear indentations by actin fibres were imaged by chromatin and actin co-staining (**Fig. 1A-B**). Using z-stack, single NPCs can be segmented in 3D based on their fluorescence intensities. Once NPCs had been segmented, we reconstructed the upper nuclear surface by interpolation (**Fig. 1C**). The punctual nature of NPCs allows a precise sampling of the NE surface by using the center of mass of their signals, contrarily to the diffuse signals of non-punctual structures stained by DAPI or lamin B1 antibody for example. When possible, we also segmented actin fibres indenting the nucleus and used the contact points in the interpolation to better reconstruct the indented area (**Fig 1C, see methods**). Reconstructed surfaces are typically not smooth but with dips and bumps. These surfaces qualitatively correspond to the non-smooth nuclear surface observed in electronic microscopy^27,28^.

Because surfaces are reconstructed from NPCs, we need to ensure that the reconstruction is robust to any missed NPCs. To this end, we applied *in silico* thinning of NPCs, i.e., we randomly excluded NPCs and measured the effect on surface reconstruction (**S1**). Thinning has a local impact on the surface reconstruction at the localization of thinned NPCs, irrespective of the thinning fraction (**S2A**). However, the global error, evaluated over the entire surface, increases with the fraction of thinned NPCs but remains reasonably low (about 30%) even with extreme thinning of 50% (**S2B**). Still, even if NPCs are numerous (878 ± 246, NPC per nucleus), they cover a small fraction of the overall surface and thus the global surface is not significantly affected by missed NPCs. In addition, we performed actin fibres segmentation to add an extra layer of precision to the reconstruction by localizing indented areas that have more complex geometry with higher curvatures. In conclusion, SIM imagining of single NPCs allows accurate reconstruction of NE.

### NPCs are not randomly distributed but dispersed at short scale

Since we were able to reconstruct the nuclear surface, we first investigated the spatial distributions of NPCs over the nuclear surface. Because the typical NPC diameter (∼100nm) is lower than distances between NPCs, we can view that data as a pattern of points on a curved surface, i.e., a Riemann manifold. If points are uniformly randomly distributed and independent of each other, the point pattern follows so-called Complete Spatial Randomness (CSR). Diggle notes two main deviations to CSR, either clustering (i.e., aggregation) or dispersion (i.e. repulsion between points forming a regular grid)^29^. To investigate if a point pattern follows CSR, the Ripley’s K function is a valuable tool^30^. It quantifies the average number of neighbors in a radius *r* around points normalized by the total point density. In cases of CSR, the Ripley K function is equal to *πr*^2^. Hence, values of K(r) > *πr*^2^ indicate more neighbors than expected, i.e., clustering of points, and conversely, values of K(r) < *πr*^2^ indicate repulsion between points. Ripley’s K function has the great advantage of quantifying spatial organization at different scales *r*, rather than summarizing the organization to a single number. However, Ripley’s K function requires knowledge of pairwise distances between NPCs. Euclidian distance is not the relevant metric, as it does not follow the nuclear surface unlike geodesic distances. However, geodesic distances are both difficult to approximate and computationally expensive, and geodesic-estimating algorithms are based on approximations^31^. Therefore, we relied on a different strategy. We projected all the points in the 2D xy plane and computed the Ripley’s K function based on classical Euclidian distances in the projected plane. As a result, areas with a higher slope tend to accumulate more points after projection, without being a genuine signal of clustering. Consequently, the Ripley’s K function computed from the projected points, is not expected to be equal to *πr*^2^ in case of CSR. To account for this projection bias, we generated CSR simulations over the same experimental surface and computed a reference Ripley’s K function from these simulated points (Fig. 2A). CSR simulations over experimental surfaces are carried out by assigning to each surface element of the reconstructed surface a probability of receiving a point directly proportional to its surface area (**see methods and Fig. S3**). Strikingly, the experimental Ripley’s K function is lower than reference CSR simulations at short scales, indicative of repulsion between NPCs. The typical length scale is ∼400nm (**Fig. 2B**). Because NPCs are typically associated with lamin fibres^32– 34^, we stained for lamin to explore whether this association could explain the observed inter-NPC repulsion. In line with previous observations, NPCs were indeed observed between lamin fibres, potentially in close apposition (**Fig. 2C**). In our C2C12 model specifically, we typically observed NPCs between lamin fibres. Hence, the typical NPC-exclusion size of 400nm could directly reflect the mesh size of the lamin network, since 400nm falls within the range of lamin mesh sizes (50-400nm) reported in the literature^34,35^. At higher scales, NPCs tend to be more randomly distributed. In conclusion, NPCs density exhibits local random fluctuations at the micrometer scale, even though NPCs have sub-micrometer repulsions.

**Figure 2.**
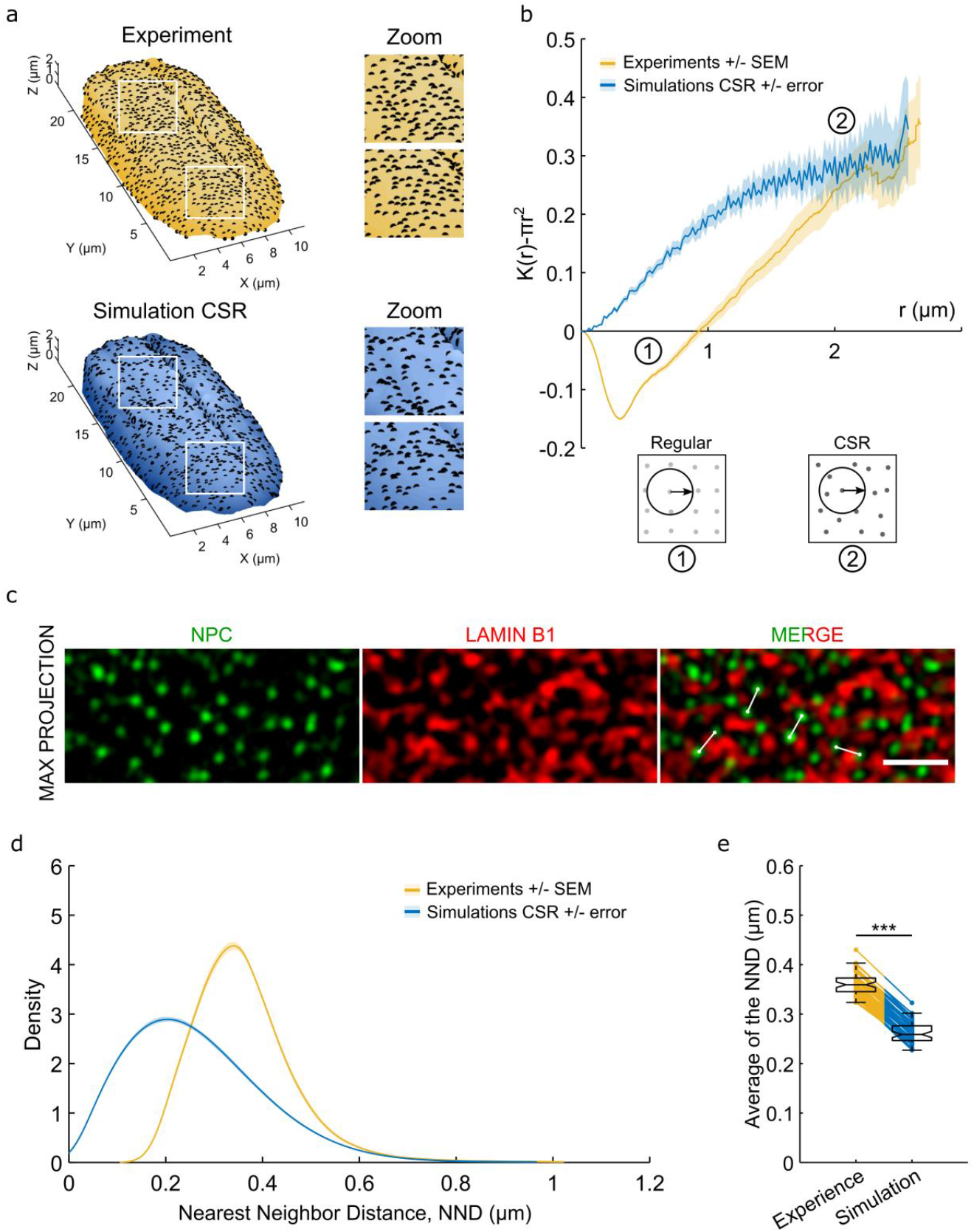
NPCs are dispersed at short scale. **a**. Experimental nuclear surface with black dots representing either experimental NPCs (blue surface) or CSR simulation (yellow surface). Both conditions have the same number of NPCs. Insights emphasize the spatial repulsions of experimental NPCs while the CSR insights present a mixture of local aggregation and repulsion. **b**. Average centered Ripley K function (*K*(*r*) − *πr*^2^) of 57 nuclei (and 50.038 NPCs) from 7 independent experiments. Yellow curve is the average of experimental data ± SEM. Blue curve is the average of 100 simulations and the blue shade is the envelope containing 95% of CSR simulations. The simulated curve is not equal to 0 due to projection effects. Hence, experimental curve should be compared to this reference. Numbers describe the type of spatial patterns observed at different scale. **c**. SIM image of lamin B1 and NPC staining. Merge shows that a lamin fibre often separates two NPCs as underscores by white lines of 370 nm. Scale bar is 1µm. **d**. Density of probability of NND from experimental data (yellow) and CSR simulations (blue). Yellow curve is the average of experimental data ± SEM. Blue curve is the average of simulations and the blue shade is the envelope containing 95% of CSR 100 simulations. **e**. Average NND comparison between experimental data and CSR simulation. Each point is a nucleus, and a single CSR simulation is realized over the same surface defining the pairing. The significance has been computed using a paired t-test. ***p<0.001. Paired Cohen’s d = −11.33).

To confirm the observed repulsion effect, we measured the Nearest Neighbor Distance (NND) of the NPCs. NNDs were measured as simple 3D Euclidian distances, reasoning that Euclidian distance is a good approximation of geodesic distances at short scales. The average probability density of observed NNDs had a mode of ∼400nm (**Fig. 2D**), confirming the NPC repulsion length estimated by the Ripley’s K function. While on flat surfaces, theoretical NND distribution under a CSR is well known, NND distribution on complex nuclear surfaces cannot be determined analytically. Therefore, we relied once again on CSR simulations on experimental nuclear surfaces. The resultant distribution has a mode of 200nm, thus confirming the existence of the observed inter-NPC repulsion (**Fig. 2D**). Even considering a steric repulsion due to NPC diameters (∼100nm), it is not sufficient to explain the 200nm shift observed between CSR simulations and experimental observations. This NPC-repulsion effect is present in almost all nuclei, as shown by the comparison, per nucleus, of observed NND with CSR NND (**Fig. 2E**). One possible explanation for this discrepancy is that the observed NPC repulsions were due to weak segmentation that interpreted several NPCs as a single NPC, thus creating an artificial repulsion. To control this possibility, we examined the relationship between segmented NPC volume and NND (**Fig S4**). Over a range of 5 times the NPCs volume (capturing ∼95% of the NPC volume, **Fig. S4A**), NND increased by only ∼100nm (**Fig. S4B)**. This control shows that this bias, if present, is far from sufficient to explain the observed repulsion.

In conclusion, our analysis shows that NPCs are not entirely randomly distributed, but instead are repelled at short scales and located more randomly at longer scales. Moreover, the typical NPC repulsion distance observed is not consistent with a simple steric repulsion of NPCs, but could reflect the additional effect of lamin meshwork.

### NPC surface density is lower in heterochromatin and indented domains

We next sought to identify the determinants of NPC localization. Upon computing the local NPC surface density, we observed clear areas of lower and higher densities (**Fig. 3A**) that seem not to be detected by the Ripley’s K function analysis as a significant deviation of CSR. As a reference, the total NPC surface density is 3.66 ± 0.6 NPC.µm^−2^ (n=57 nuclei) (**Fig. 3B**) corresponding to a number of NPCs on the upper nuclear surface of 878 ± 246 and an upper nuclear surface of 247 ± 83µm^2^. However, we explored this density heterogeneity with a more direct statistical comparison by looking for the signature of NPCs spatial segregations. In particular, cells spread on elongated adhesive micropatterns show apical stress fibres indenting the nucleus (**Fig. 1A-C**). These indented regions seem to exhibit a lower NPC surface density, although other low density non-indented regions also exist (**Fig. 3A**). On the basis of actin staining (with SiR-actin), we segmented indented regions (**Fig. 3C**). When possible, we compared the NPC surface density assessed on indented and non-indented areas of the same nucleus (**Fig. 3D**). Such a pairing per nucleus excludes any potential staining bias between nuclei. We found that indented areas had a lower NPC density (3.35 ± 0.88 NPC.µm^−2^ vs 4.02 ± 0.71 NPC.µm^−2^, n=14 nuclei). Our observations contrast with those of the Beckerle’s group who reported a higher NPC density on indented areas^17,18^. This discrepancy could simply be explained by the possibility that actin fibres also indent the nucleus in Beckerle’s group’s model, but that this indentation was not detected due to limited optical resolution. As this indentation generates curvature, it increases the membrane area imaged which can then be interpreted as NPC signal enrichment when projected onto the x-y plane, without any real increase in NPC surface density.

**Figure 3.**
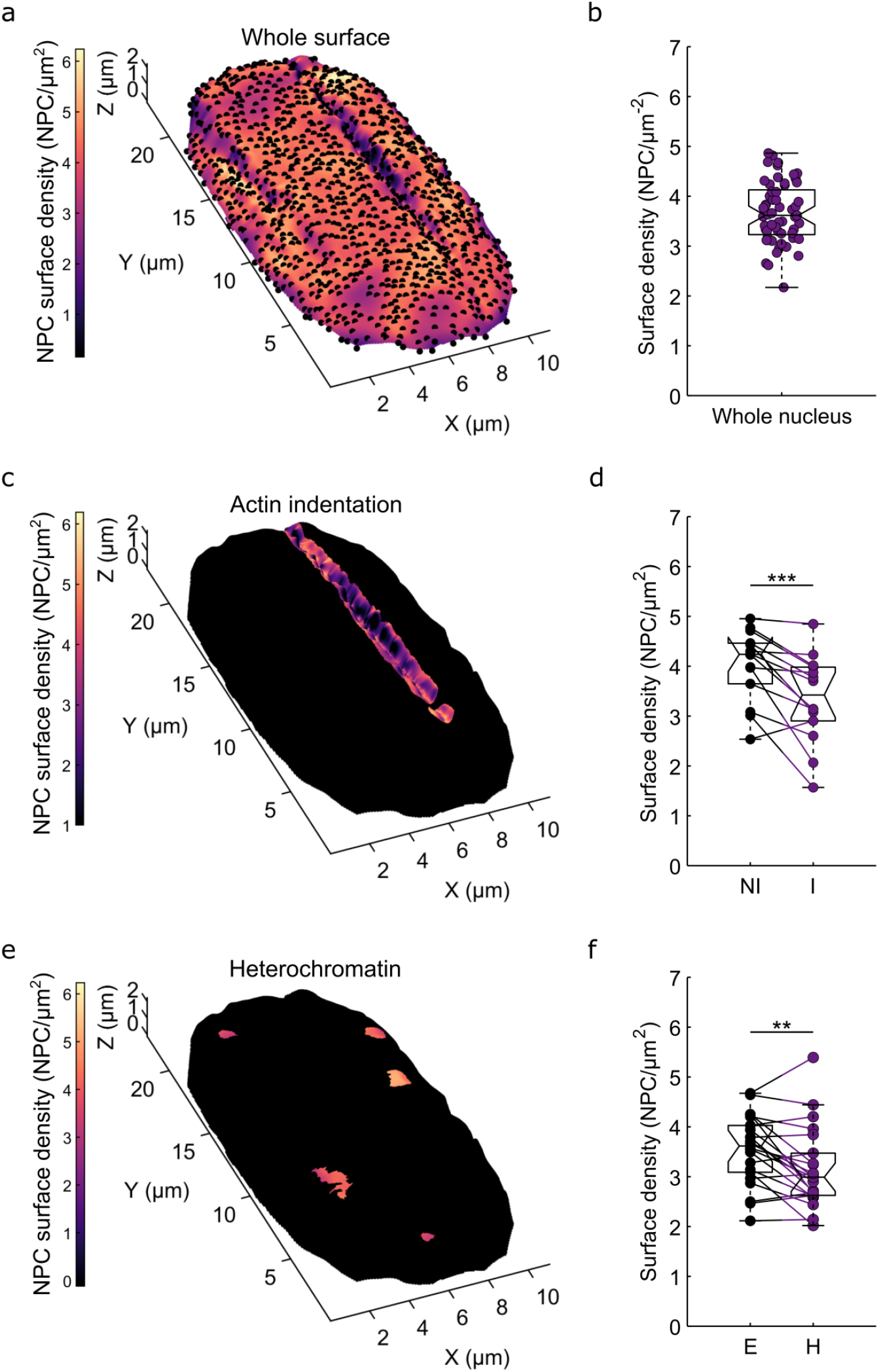
NPC density correlates with chromatin state and indentation. **a**. Nuclear surface with black dots representing NPCs and color code represents the local NPC surface density. **b**. Boxplot of total surface density per nucleus simply computed as the total number of NPC divided by the nuclear surface. NPC surface density 3.66 ± 0.6 NPC.µm^−2^ (evaluated o 57 nuclei from 7 independent experiments). **c**. Same nuclear surface with NPC surface density represented only on the indentation area. **d**. Comparison of NPC surface density on non-indented and indented areas: 3.35 ± 0.88 NPC.µm^−2^ vs 4.02 ± 0.71 NPC.µm^−2^ (evaluated on 14 nuclei from 3 independent experiments). Paired Cohen’s d = −1.09. **e**. Same nuclear surface with NPC surface density represented only on heterochromatin domains. **f**. Comparison of NPC surface density on euchromatin and heterochromatin domains: 3.15 ± 0.80 NPC.µm^−2^ vs 3.54 ± 0.69 NPC.µm^−2^ (evaluated on 22 nuclei from 3 independent experiments). Paired Cohen’s d = −0.70. In d and f, each pair of points is a nucleus divided in the two aforementioned regions. Surface densities are computed as the number of NPCs in the region of interest divided by its surface. The significance has been computed using a paired Wilcoxon test. **p<0.01 and ***p<0.001.

Heterochromatin domains could be another candidate for the signature of NPCs spatial segregation. We hypothesized that regions of the NE near to large heterochromatin domains-viewed as inactive chromatin-would have lower NPC density (**Fig. 3C**). On the basis of chromatin staining (by DAPI), we therefore segmented heterochromatin domains (**Fig. 3E**). Once again, the pairing per nucleus excludes any potential staining bias between nuclei. The analysis of heterochromatin domains also revealed lower NPC surface density in these regions (3.15 ± 0.80 NPC.µm^−2^ vs 3.54 ± 0.69 NPC.µm^−2^, n=22 nuclei) (**Fig. 3F**).

In conclusion, our results shed light on two correlates of NPC localization: chromatin and mechanical states.

### NPC density correlates with local membrane curvature

In the previous section, we show that the NPCs surface density correlates with the chromatin and the mechanical states. An additional physical element that could affect the NPCs spatial distribution is the curvature of the NE. Indeed, some of the NPC’s components are believed to be sensitive to membrane curvature^20^. In order to test the potential effect of curvature on NPCs spatial distribution we relied on bi-harmonic spline interpolation and the Laplace-Beltrami operator to generate curvature maps of nuclear surfaces (**Fig. 4A, C, methods and SM**). It is important to note that, our previous thinning analysis revealed that missed NPCs can generate local variation in the surface area (**Fig. S1 and S2A**). We extended the thinning analysis to curvature and indeed found that missed NPCs can generate strong error on local curvatures (**Fig. S5 and S6**). However, this error is extremely local and can be mitigated by evaluating curvature not on punctual locations but by region (**S2C-E**). Therefore, we applied a binning on the curvature maps in our data to ensure robustness to local errors, although this reduced resolution. Next, we plotted the local NPC surface density as a function of Gaussian (K) and mean (H) curvatures (**Fig. 4B, 4D**). We observed that the NPC surface density-Gaussian curvature relationship exhibits a slightly asymmetric bell curve centered on K=0 (**Fig. 4C**). This preference for K=0 could potentially reflect mechanical constraints that make it difficult for the membrane to bend in two directions around a NPC (saddle point). Similarly, the NPC surface density as a function of mean curvature is also an asymmetric bell curve but centered on H=−0.23 µm^−1^ (Gaussian fit) (**Fig. 4D**).

**Figure 4.**
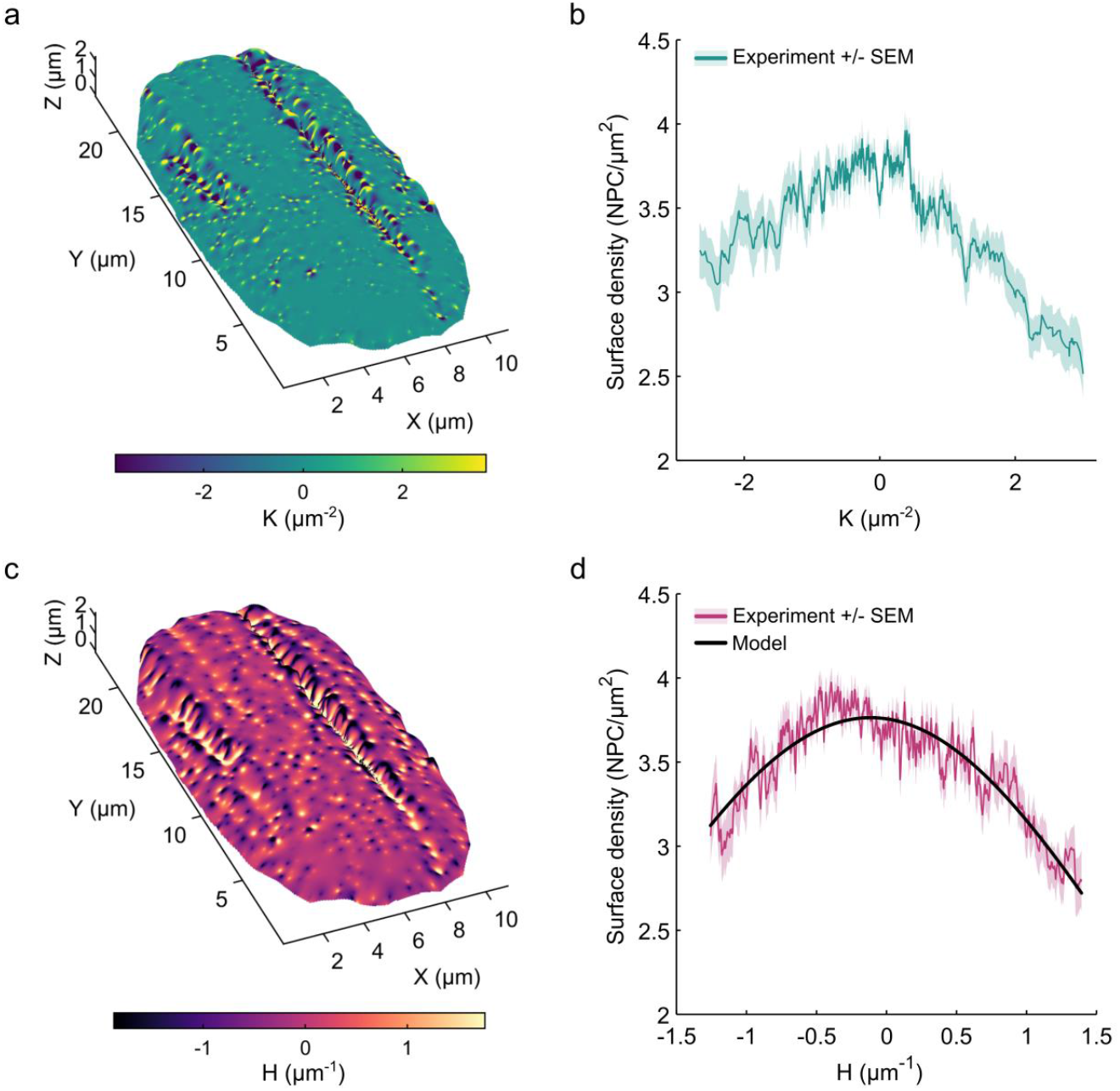
NPC density correlates with membrane curvature. **a**. Nuclear surface with color code representing the local Gaussian curvature K. **b**. NPC surface density as a function of the Gaussian curvature K. Solid line represents the average and the shade the SEM. **c**. Nuclear surface with color code representing the local mean curvature H. **d**. NPC surface density as a function of the mean curvature H. Orange solid line represents the average and the shade the SEM. Black solid line represents the fit with the theoretical model. In b and d, data are coming from 57 nuclei from 3 independent experiments. In b and d, a 16×16 (640×640nm) binning is applied on the NPC surface density and curvatures to be robust against local errors and measure curvature sensing at a scale bigger than the NPC size.

To ensure that these observed curves reflect actual relationships between NPC density and curvatures, and not artifacts of the method, we simulated different nucleus geometries (flat, hemi-spherical or ellipsoidal) (**Fig. S7A and S8A**). We generated random distributions of NPCs on these surfaces with the same surface density as observed in the experimental data. Of note, the simulated NPCs are not CSRs *stricto sensu* because we added a 100nm repulsion length between NPCs to model their spatial expansion and to better match the experimental data. Such a pattern of dots is formally defined as a Matérn II hard-core process. Furthermore, we also added noise to simulated NPCs in the z direction to incorporate uncertainty on the z position of NPCs as well as to generate fluctuations like the experimental NE surfaces which are not smooth (**Fig. S7A and S8A**). We reconstructed the surfaces from simulated NPCs and computed the associated curvature using the same method as previously described. We then explored the relationships between NPC surface density and Gaussian and mean curvatures on these synthetic data (**Fig. S7B and S8B**). Over the range of simulated geometries and noises, we did not reproduce our experimental data. Because surface density is constant over the surfaces, we do not expect any correlation between density and curvature on these synthetic data. However, the addition of noise can create correlations that are therefore due to the methods. We still argue that the bias introduced by the method cannot explain our observations. Instead, the potential bias seems to be the other way around, creating an V-shaped curve and not a bell curve as observed. Finally, we extended the thinning analysis to the density-curvature relationship (**Fig. S9A-B**). We found that removing up to 20% of the NPCs does not affect the shape of the curves. Altogether, these results and extensive simulations controls indicate that this correlation is genuine and not the result of a bias introduced by the method.

In conclusion, Laplace-Beltrami operator is a robust method to estimate curvatures on discrete surfaces. Moreover, we observed a non-monotonous relationship between NPC density and Gaussian and mean curvature. Extensive simulations controls seem to indicate that this correlation is genuine and not the result of a bias introduced by the method.

We thus observed a depletion in NPC for highly positively or negatively curved regions of the NE.

### The relationship between mean curvature and NPC surface density is compatible with the existence of a NPC preferential curvature

Finally, we wondered whether the existence of a preferential curvature of the NPC could explain the observed relationship between density and curvature (**Fig. 4D**). To better understand our results, we compared our experimental observations with a statistical physics modeling. We applied a model close to that of Yang et al.^36^, which is similar to previous theoretical approaches^37,38^. We hypothesize that the energy associated with the insertion of a NPC in the NE contains a curvature-sensitive term:

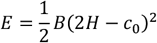

B being an effective bending modulus of the complex NE-NPC, and c_0_ the preferential curvature of the NPC. The Gaussian curvature distribution is constant according to the Gauss-Bonnet theorem and can thus be ignored. The NPC surface density *φ* will then follow a Botlzmann distribution:

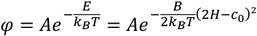

k_B_T being the thermal energy and A the NPC surface density when *H* = *c*_0_. This analytical model captures well the observed distribution (**Fig. 4F**). The preferred curvature c_0_ is −0.23 µm^−1^ (Bootstrap 95% confidence interval [-0.326; −0.114]) which corresponds to a local radius of curvature *R*_0_=1/*c*_*0*_= −4.35 µm. Note that the value of the optimal radius of curvature is similar to the radius of curvature of the nucleus before cell spreading: 5.14 ± 0.59 µm, calculated as the radius of a sphere with a volume equal to the one we measured for the nuclear volume of spread cells, 591 ± 217 µm^3^. Of note, the measurement of the volume through the NPC reconstruction is a good way to avoid biases on volume calculation due to optical aberration. Curvatures below and above *c*_*0*_ lead to lower values of NPC density. Our data indicates a coupling between local membrane curvature and NPC density. We propose as a working hypothesis that this coupling potentially roots from the mechanics of the NPC and the existence of a preferred curvature.

### Conclusion and discussion

In this study, we combined micropatterning technique with super resolution microscopy (SIM) to image NPCs and reconstruct the 3D nuclear surfaces. Spatial analysis of NPC distribution, using Ripley’s K function, simulations and NND analysis revealed that NPCs are not entirely randomly distributed. Instead, they exhibit short-range repulsion and more randomly located at longer scales. Although Ripley’s K function is limited to ∼2µm to minimize borders effects, however, we cannot exclude the possibility that NPCs also cluster at higher scales, as suggested by the high slope of Ripley’s K function at ∼2µm. Moreover, the typical repulsion distance of ∼400nm is not compatible with a simple steric repulsion of NPCs, indicating that the lamin meshwork may contribute to this spatial arrangement.

We also observed that actin-indented areas of the nucleus have a lower NPC density, a finding that contrast with reports by Beckerle’s group, who found higher NPC density on indented areas^17,18^. This discrepancy may be simply explained by the limited optical resolution of the microscope (laser scanning microscope) used or due to a difference in cell type (mouse fibroblast vs myoblast) resulting in NPCs with different inherent molecular composition and stoichiometry^39,40^. Additionally, Beckerle’s group also observed that NPC staining near actin bundles is, which could create the appearance of NPC enrichment^17^. However, the origin of this modulation in fluorescent brightness remains unclear. Another potential explanation for this discrepancy is that the actin lines observed in fibroblasts do not indent the NE, as suggested by Beckerle’s group, and therefore represent distinct structures from those seen in C2C12 mybolasts^17^. Alternatively, actin fibres in Beckerle’s group’s model may actualle indent the nucleus, but this could have been overlooked due to limited resolution. If such indentation occurs, it generates curvature, increasing the imaged membrane area, which could lead to an artifact where the NPC signal appears enriched when projected onto the x-y plane, despite no true increase in NPC density. This highlights the importance of accurately considering the correct NE geometry when assessing NPC distribution in indented regions.

We also observed that NPC density is lower in heterochromatin domains, an interesting finding in the context of the newly discovered roles of NPCs in gene regulation and genome organization^41–43^. While, the interaction between NPCs and the thin layer of heterochromatin upholstering the internal membrane of NE is well studied, the spatial correlation between NPCs and larger heterochromatin regions has been less characterized. Further investigations are required to elucidate whether these spatial correlations are causative relationships and if they support any functional coupling. While it may seem intuitive that fewer NPCs are located near inactive chromatin, the reduced NOC density in indented areas is more difficult to explain. One possibility may be that the lower NPC density in these regions is simply a mechanical consequence of the indentation, which stretches the NE while maintaining a constant number of NPCs.

Previous studies have highlighted the role of membrane curvature in protein sorting^36–38^. Our measurements reveal a correlation between local curvature of the NE and the recruitment of the NPC. We demonstrate that NPC density is influenced by the geometry of the NE and we determine the most favorable curvature for this complex *c*_*0*_=−0.23 µm^−1^. The interaction between NPC density and NE curvature may be mediated by the presence of curved motives or amphipatic helices in the NPC, that operate as curvature sensors. Notably, the arrangement of the Y complex (part of the NPC) has been compared to the structure of coatomers ^22,23,44^. Nup133, which contains the curvature-sensing ALPS domain, is particularly promising, as its binding to liposomes is dependent on their radius of curvature^45^. However, since mammalian NPCs have limited mobility, if any^5^, our findings are more likely to reflect a preference for a specific curvature c_0_ during NPC biogenesis and especially during their insertion into the membranes. In addition, it has been recently suggested that lamins are diluted in regions of high Gaussian curvature^6^. This dilution could hypothetically reduce the interaction between NPCs and lamins, and consequently increase NPCs mobility.

In conclusion, by combining micropatterning technique, super resolution microscopy, surface reconstruction and spatial statistics, we were able to show a specific spatial distribution of NPCs and a correlation between NPC density and NE curvature. Our study represents the first step toward understanding the role of curvature on NPCs density, paving the way for a new description of the properties of NPCs.

## Acknowledgments

We acknowledge the ImagoSeine core facility of the Institut Jacques Monod (member of the France BioImaging, 566 ANR-10-INBS-04). The authors are grateful to Caitlin Martin for her diligent proofreading of the manuscript.

## Author Contributions

DP and HL contributed equally to this work.

DP, SH and FM developed the conceptual framework and designed the study. SH sought funding. DP performed all experiments and supervised the study. HL performed all quantitative analysis and simulations. OF and FM helped with experiments.

DP, HL, OF, FM and SH analysed the findings and wrote the manuscript. All the authors reviewed, edited and approved the paper.

## Conflict of Interest

*The authors declare that the research was conducted in the absence of any commercial or financial relationships that could be construed as a potential conflict of interest*.

## Funding

This work was supported by the LabEx “Who Am I?” #ANR-11-LABX-0071 567 and the Université de Paris IdEx #ANR-18-IDEX-0001 funded by the French Government 568 through its “Investments for the Future” program.

## Methods

### Segmentation

#### Segmentation of NPCs

The function “3D object counter” of ImageJ was used to segment NPCs. The intensity threshold has been manually adjusted for each image. Aspecific WGA staining from the membrane or the cytosol is sometimes present in close apposition of the nucleus. To retain only the NPC, an intensity-based segmentation of the nucleus from lamin or DAPI staining is realized. The nucleus ROI is then set to 300nm and NPC segmentation is only performed on this ROI.

#### Segmentation of nucleus and chromatin

For images with DAPI or a lamin staining, nucleus intensity-based segmentations with manual thresholding were done after a max intensity z-projection. Holes in the mask and structures outside of the nucleus masks were automatically removed with a custom R code. Moreover, images with DAPI staining were segmented for the apical heterochromatin following the same methods (z-projection is applied only on the most apical slices).

#### Segmentation of actin Bundles

When the nucleus is indented by a SiR-actin stained actin fibre, the latter is segmented. First, a 3D ROI is manually drawn to isolate the fibre before an intensity-based segmentation with manual thresholding is done. The segmentation is performed on the x-z or y-z plane and coordinates of the centroid of each actin fibre are extracted for each plane.

### Actin Bundles Reconstruction

After segmentation, the actin bundle is a list of centroid points. To reconstruct the bundle, first, points outside nucleus masks are excluded. Second, clusters of points are determined with the DB-SCAN algorithm (Rpackages fpc and dbscan)^48^. Aberrant clusters (i.e. group of points from aspecific staining around the fibre) are manually removed. As a result, we obtain a group of points forming the skeleton of the actin bundles.

Next, we replace each of these “skeleton” points by 5 points at the surfaces of the actin bundle in contact to the nuclear envelope. These points are generated through 4 successive steps for each skeleton point:

1) We identify the first and second neighbors in the x-y plane of the skeleton point of interest.
2) We determine the line crossing the skeleton point of interest with the same slope as the segment crossing the 2 neighboring skeleton points. This line represents the local direction of the bundle.
3) 5 secondary skeleton points are created on this line at distances of −20, 10, 0, −10 and 20nm respectively from the primary skeleton point of interest.
4) Final points are computed at the surface of a cylinder with a diameter of 100nm and centered on the 5 secondary skeleton points. The position of the cylinder to these points is determined by arbitrary angles (2.8*π*/2, 2.9*π*/2, 3*π*/2, 3.1*π*/2, 3.2*π*/2) that forces contact with the nuclear envelope (i.e. the points are below the fibre, in the region of nucleus contact). Each of the 5 secondary skeleton points are randomly linked to one angle generating the final actin points.

### Surface Interpolation

The interpolation is computed from NPC centroids in addition to the actin points (see *Actin Bundle Reconstruction*) once an actin bundle has been segmented. First basal NPCs are filtered thanks to a criterion on the z coordinate. Aberrant NPC points outside the nucleus mask or inside the nucleoplasm are also manually filtered.

Next, an initial surface interpolation conducted only using NPC points is realized with the function griddata of MATLAB. We set the step size equal to the pixel size, i.e., 39.7nm. We applied a cubic interpolation or a biharmonic spline interpolation (so-called v4 by MATLAB) and we compared performances. Both interpolations generate class C^2^ surfaces but while the cubic interpolation is based on triangulation, the biharmonic is not, and hence avoids some artifacts (**see Fig. S10B and SM**).

Following initial interpolation, a manual quality control is performed on each nucleus, and artifactual NPC points generating spurious discontinuities in the surfaces are removed. Nuclei with initial NPC segmentation insufficient for good surface interpolation are filtered at this step. Similarly, actin points that too far (>500nm) from the computed surfaces are also filtered. A second interpolation is performed on the remaining NPC and actin points as described above.

The v4 interpolation generates artefacts on the border of the rectangular grid where no NPCs are detected. We computed a convex hull (R function chull) in the x-y plane on NPC coordinates and applied it as a mask to remove these artefacts.

Following this process, our surface is numerically represented as a matrix where column and row indices give the x and y coordinates while the values of the matrix give the surface height h.

### Curvature

To compute curvatures, we compared the performances of two methods:

#### Finite Difference Method (FDM)

The surface in a Monge gauge is described as the height of the surface *h* for any (*x, y*) couple, hence *h* = *f*(*x, y*). In Monge gauge, principal curvatures *k*_1_ and *k*_2_ are the eigenvalues of the Hessian matrix of the surface Λ ^49,50^:

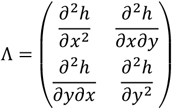

The partial derivatives are tractable since the two interpolation methods used generate C^2^ class surfaces. The derivatives can be numerically calculated by FDM, i.e., partial derivative operators are computed by the convolution of a matrix with the surface *h*. Hence:

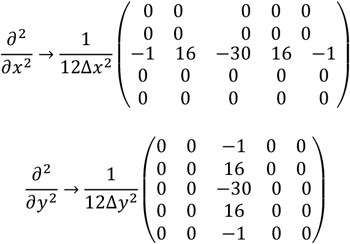

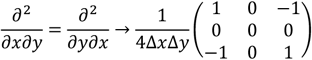

With Δ*x* = Δ*y* the step size i.e. the pixel size^51^. Then, eigenvalues can be easily computed from the hessian matrix. Once principal curvatures are computed, we compute mean and Gaussian curvatures respectively as:

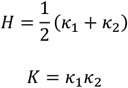

#### Discrete Laplace-Beltrami Operator

The Laplace-Beltrami operator is a generalization of the Laplace operator for Riemannian manifolds such as smooth surfaces. In this context, the Laplace-Beltrami operator of the surface is the trace of the Hessian matrix, *i*.*e*., half of the mean curvature. Hence, this method to compute the curvature is mathematically equivalent to the one based on the eigenvalues of the Hessian. However, as our numerical data are discrete, we need to refer to the discrete Laplace-Beltrami operator (**Fig. S10A**), resulting in different approximations and results from the FDM computation of the Hessian. We followed the method of Meyer et al. ^52^. The nuclear surface is described as a triangular mesh. Let *v* be one of the vertices, *N*(*v*) the set of neighbors associated to the vertex *v*, and 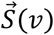 a vector-valued function giving coordinates of the vertex *v*. For such a triangular mesh, the discretization of Laplace-Beltrami operator leads to the so-called cotangent formula:

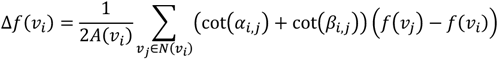

Where *A*(*v*_*i*_) is the area associated to the vertex *v*_*i*_. As is classically done, this area is computed as the “barycentric cell area” i.e. one third of the sum triangles’ area in which vertex *v*_*i*_ is involved. The angles *α*_*i,j*_ and *β*_*i,j*_ are the opposite angles of the two triangles in which the edge between *v*_*i*_ and *v*_*j*_ is involved. This cotangent term is called the weight. *f*(*v*) is a function defined on the triangular mesh. The application of the Laplace-Beltrami to the function 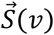 gives the mean curvature:

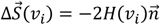

With *H* the mean curvature and 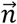 the surface normal vector.

Additionally, the Gaussian curvature *K* of such a triangular mesh can be computed according to the angle deficit method^53^:

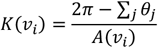

For the triangle *j* in which *v*_*i*_ is a vertex, *θ*_*j*_ is the angle formed by the two triangle edges crossing at *v*_*i*_. From Gaussian and mean curvature, principal curvatures can be computed as:

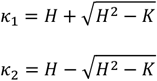

The triangular mesh describing nuclear surface has been computed first using a Delaunay triangulation in the x-y plane. Because our grid is regular i.e. nodes are evenly spaced at a distance equal to the pixel size, the Delaunay triangulation is trivial and computationally cheap. However, adding the height of the point leads to some triangle edges that are not longer Delaunay. This can then produce negative weights in the computation of the Laplace-Beltrami operator. In our date, therefore, we checked that these negative weights typically represent <1% of nuclear surfaces. Moreover, computation leading to *H*^2^ − *K* < 0 typically represents <0.1% of the nuclear surfaces.

#### Comparisons of the methods

To evaluate numerical robustness of the two methods (i.e. FDM and discrete Laplace-Beltrami Operator), we generated synthetic hemispherical surfaces of 1µm with different pixel sizes (i.e. different sampling depth) and computed the curvatures with the two methods. On such geometry, mean, Gaussian, and principal curvatures are all equal to −1µm^−1^, therefore we restrict our analysis to the principal curvature *κ*_1_. For each position in the hemisphere, we compare the theoretical curvature to the computed one. The error is defined as 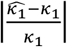. It is clear that FDM generates non-constant curvatures on the sphere surface, as the higher the derivatives (according to x or y), the higher the curvature and the error. On the other hand, the Laplace-Beltrami approach generates almost constant curvatures across the surface, with the exception of the extreme vertices. The higher the resolution of the hemisphere (i.e. the higher is the sampling of the surface), the lower is the median error. However, the tail of error distribution becomes longer while less important in proportion; in other words, the error tail becomes more extreme but concentrated on fewer vertices at the surface borders (**Table S1**).

### Spatial statistics

#### Density

NPCs can be seen as a point pattern lying on a Riemannian manifold *i*.*e*. a surface. This kind of point pattern is analyzed by spatial statistics, where the first property of a point pattern is defined as its intensity *λ*, that can be interpreted as a local surface density:

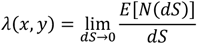

where *x* and *y* are the coordinates in a Monge gauge, *dS* the surface element, and *N*(*dS*) the number of points in the surface element centered at (*x, y*). The intensity is easy to compute for a point pattern lying on a 2D Euclidian space (i.e. a plane) but more complicated to evaluate on a Riemannian manifold. We follow the method developed in Baddeley et al.^54^. Briefly, Let *λ*_*p*_ be the intensity of the point pattern relative to the 2D projected plane surface *P* and *λ*_*s*_ be the intensity relative to the actual surface *S*. We have the following relationship:

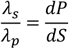

With *dP* representing the surface element in the projected plane and *dS* the surface element on the Riemannian manifold. Considering that the projected surface element *dP* corresponds to a single pixel in the Monge gauge of the surface with side length Δ*x* = Δ*y*, we have:

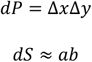

With *a* and *b* the length side of the surface element *dS* (**Fig. S3A**). Note that the surface element *dS* is not necessarily planar and therefore we use a first order approximation for its area. From these relationships we can deduce the following:

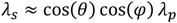

With *θ* and *φ* the angles between the *x* and *y* axis and the sides of the surface element *dS* (**Fig. S3A**).

One can rewrite the equality as:

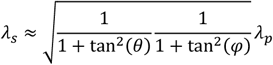

And approximate the tangents as derivatives:

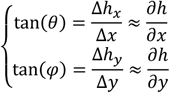

With *Δh*_*x*_ and *Δh*_*y*_ the lengths of the triangle sides opposite to angles *θ* and *φ* respectively (**Fig. S3A**). Hence, the expression of the point pattern intensity becomes:

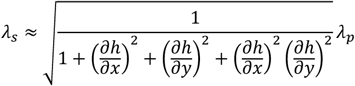

The derivatives can be numerically calculated by FDM i.e. partial derivatives operators can be computed as the convolution of a matrix with the surface *h* ^51^:

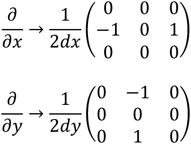

Hence, the projected point pattern intensity if first calculated with the function density from the R package spatstat^54^ with a Gaussian kernel using the Diggle’s method for bandwidth selection^55^. Then, the derivatives using FDM are computed as explained above to convert *λ*_*p*_ into *λ*_*s*_. Note this method also allows estimation of nucleus element surface *dS* and total surface *S*:

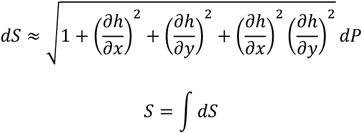

#### Ripley’s K function

For a set of *n* NPC coordinates *X* = {*x*_1_, *x*_2_, *…*, *x*_*n*_} observed in an area |*Ω*|, The Ripley’s K function^56^ is defined as:

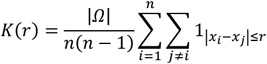

This function quantifies the average number of NPC in a disk of radius *r* centered on one NPC. In case of Complete Spatial Randomness (CSR) *i*.*e*. uniformly and independent distribution of points, the estimator of this function should fluctuates around *πr*^2^. Therefore we subtracted *πr*^2^ to *K*(*r*). Hence, a positive value indicates clustering whereas a negative value indicates dispersing/repulsion.

The direct computation on the Riemannian manifold is complicated because it requires to compute geodesic distances between NPCs on the surface. We simplified the problem by computing the Ripley’s K function on NPC projected on the x-y plane. We used the function Kest of the R package spatstat with the isotropic correction for boundary effects. We used the Ripley-Rasson estimate of the spatial domain that produced better result than a simple convex hull^57^.

However, clustering can be observed because of the projection i.e. points in region with a high slope tend to be closer in the projected version. To control this effect, instead of just comparing the Ripley’s K function to the theoretical expectation we run Monte-Carlo simulations (see *Monte-Carlo Simulations section*) on the same surface, projected these simulations and computed the Ripley’s K function. This approach gives the reference for a CSR pattern projected from the actual surface.

Hence, Monte Carlo simulations were used to generate CSR envelopes as in Lachuer et al^58^. Because observed Ripley’s K function is averaged over a population of nucleus; CSR simulations were run in the same way. For each nucleus, a Monte Carlo CSR simulation is run using the observed nucleus surface and the observed number of NPCs. The resulting Ripley’s K functions are averaged over the nucleus. This procedure was repeated a high number of times (*n* = 100). The CSR envelope contains 95% of these averaged (over the nucleus) Ripley’s K functions and the solid line is the average (over the simulations). This procedure controls accurately CSR deviation to *πr*^2^ that could be due to the projection and the border effects.

#### NND

As an alternative way to quantify spatial structures of NPCs, we also measure the Nearest Neighbor Distance (NND). Still, because geodesic distances are complicated to compute on the surface, we computed the NND as the 3D Euclidian distance between NPCs irrespective of the surface. We reasoned that at small scales, 3D Euclidian distance is a good approximation of the geodesic distance. To control any possible bias induced by our approximation, we compared these experimental NND to Monte-Carlo simulations over the same surfaces as made for the Ripley’s K function.

### Monte-Carlo Simulations

#### Experimental surfaces

To simulate CSR on Riemannian manifold, the probability to draw a point in a surface element *dS* over the total area *S* is just *dS*/*S*. For nucleus surfaces, we computed all the surface elements *dS* and the total surfaces *S* (see *Spatial Statistics* section) to compute the density of probability over the surface and then draw points from it.

#### Square

To generate CSR on a square of side length *L*, we draw *x* and *y* Cartesian coordinates in uniform law (*x∼U(*0, *L)* and *y∼U(*0, *L)*) and set *z* = 0.

#### Hemi-sphere

To generate CSR on a hemi-sphere of radius *R* we draw *u* and *v* in uniform law (*u∼U(*0,1) and *v∼U(*0.*5*,1)) and obtained the polar coordinates by the following transformation:

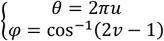

We cannot directly draw *θ* and *φ* from the uniform law because on a sphere the surface element is a function of *θ* (*dS* = *R*^*2*^ *sin(θ*)*dθdφ*). We therefore convert these polar coordinates into Cartesian coordinates:

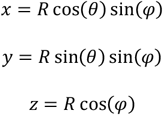

This method is already widely used^59,60^.

#### Hemi-ellipsoid

The generation of CSR on a hemi-ellipsoid is less evident. The strategy is to generate a CSR on a sphere and then deform it into an ellipsoid following the logic of the work of “Random selection of points distributed on curved surfaces” and “Simulation studies of a phenomenological model for elongated virus capsid formation” ^60,61^. However, the deformation is not uniform and the probability to observe a point in a surface element needs to be weighted by the strength of the deformation of this surface element. Let *a, b* and *c* be the lengths of the semi-axis of the ellipsoid. The equation of an ellipsoid is:

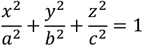

We can deform a sphere of unit radius into an ellipsoid with the following function:

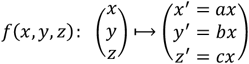

We can generate CSR on s sphere and then deform it to obtain an ellipsoid. However, a surface element *dS* on the sphere is deformed into a surface element *dS′* on the ellipsoid. Let *μ* be the multiplicative coefficient:

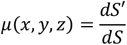

Let *M* be a point on the sphere centered on *dS* and 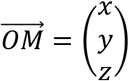 be the position vector. Let 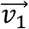 and 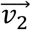 be a pair of tangent vectors to the point *M* i.e. satisfying:

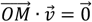

*H*ence, the following choice is valid:

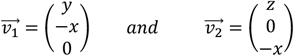

We do not need 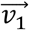 and 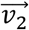 to be orthogonal to define a surface element:

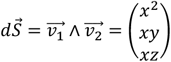

Hence:

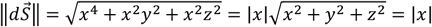

After deformation into an ellipsoid:

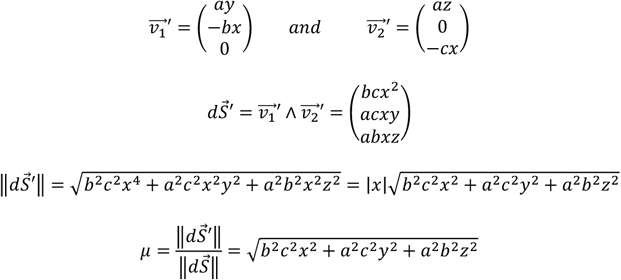

The extremums of the coefficient *μ* are at the poles of the sphere:

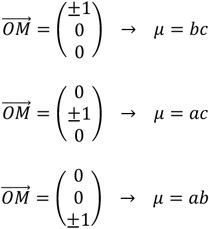

Let *μ*_*max*_ be:

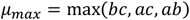

*T*herefore, the CSR on ellipsoid is achieved in 3 steps:

i) Draw points on a unit sphere in a CSR manner.
ii) Conserve each point with a probability *μ(x, y, z)*/*μ*_*max*_.
iii) Apply the ellipsoid deformation. The ellipsoid CSR can be easily transformed into hemi-ellipsoid by removing all the points *z* < 0.

### Nucleus simulations for simple geometry

To control if observed correlation with curvature and spatial repulsion could be due only to the geometry and our methods, we simulate nuclei with characteristic shapes (planar, hemi-spherical and hemi-ellipsoidal) and a random distribution of NPCs over the surface. To take into consideration the steric repulsion between NPCs, we did not model NPCs distribution over the surface as a CSR but as a Matérn II hard core process^54^. We first generated successive CSR “proposal” points. If a new CSR “proposal” point has a neighbor in a distance inferior to *r*_*repulsion*_ the point is not retained. Because two NPCs cannot be closer than 2 times their radius, we choose *r*_*repulsion*_ = 100*nm*. See the *Monte-Carlo Simulations* section for the CSR part of the simulation over the aforementioned geometries.

In addition, we add a noise on the *z*_*Matérn*_ coordinates of the NPCs simulated with the Matérn II hard core process. Hence the final z coordinate is given by *z*_*final*_ = *z*_*Matérn*_ *+ ε* in which *ε* is a Gaussian random noise. This noise modelled our lower resolution in z (110nm) as well as a local deformation of the geometry to be closer to actual surfaces. We choose three levels of noise defined by the standard-deviation of the Gaussian: 0nm, 110nm and 330nm. Moreover, in these simulations, the surface of the geometry is equivalent to the experimental measures (242µm^2^ ± 82µm^2^) and the number of NPCs equivalent to experimental measures (879 ± 244).

### Chromatin analysis

We segmented apical heterochromatin (see *Segmentation* section). The projection of these masks on the interpolated nuclear surfaces defined the heterochromatin and the euchromatin regions. We computed the surfaces of the two regions (see *Spatial statistics* section) and counted the number of NPCs to obtain the average surface density. Based on heterochromatin and euchromatin masks, we also compared the median curvatures between the two regions.

### Volume estimation

Nuclear volume has been estimated by approximating the nucleus as half of an ellipsoid based on our microscopic observation. For each nucleus, we fit an ellipse in the xy, xz and yz planes to estimate the axes lengths of the ellipsoid.

### Indented regions analysis

The analysis of the actin-indented region is similar to the one for chromatin (see *Chromatin analysis section*). To create the mask of the indented and non-indented regions, we used the coordinates of actin points (see *Actin Bundles Reconstruction section*). Around each actin point, we create a disk (in the projected x-y plane) with a radius of 15 pixels (about 600nm). The sum of all these disks defines the indented regions and the non-indented region is defined as the negative.

### Statistical analysis

All our analysis was made in R 4.2.2. The number of nuclei and the number of independent trials is indicated in the legend of the figures. Since our cells are isolated on their micro-pattern when imaged, we performed single cells analysis and each nucleus was considered as independent, setting the sample size. The statistical test used is indicated in the legend. Tests are conducted in a two-sided manner and in a paired-way because we compared different regions coming from the same nucleus. When *n* ≥ 30, we used paired t-tests and when *n* < 30 we used paired Wilcoxon’s W test. Effect sizes are reported in the form of Cohen’s d for paired data, i.e., the means difference divided by the standard-deviation of the paired differences.

### Thinning

Because the surface interpolation relies on NPC segmentation, we tested how robust is our method is to segmentations missing some NPCs. Our approach relies on NPC thinning i.e. the random elimination of a given proportion of the experimental NPCs. We tested different proportions of thinning: 2.5%, 5%, 10%, 20%, 30%, 40% and 50%. From these thinned datasets, we applied v4 interpolation to reconstruct nuclear surfaces (see *Surface Interpolation* section). We compared the surface height of the thinned nucleus *h*_*thinned*_ with the reference nuclear surface computed with all the points *h*_*reference*_. The error is computed as Δ*z(x, y*) = *h*_*thinned*_(*x, y*) − *h*_*reference*_*(x, y*). We computed the median of the |*Δ*| per nucleus over the full surface (**Fig. S1, S2A**) and only at thinned NPCs (**Fig. S2B**). Total error demonstrates that the global surface is almost not impacted by thinning even at a high proportion; while the error at thinned NPC proportions demonstrates that missing NPCs can lead to a less negligible error, irrespective of thinning proportion. We applied the same strategy to measure error on curvatures (**Fig. S2C-E**). These results demonstrate than even low error on the surface height can lead to high local error on curvatures justifying the binning approach used.

### Correlation curvature-NPC density

*Experimental correlation*. Because our thinning analysis revealed that local error is non-negligible for the curvature, we applied a 16×16 pixels binning (about 635×635nm) on the curvatures and the local density of NPCs. For each nucleus, we correlate these two quantities and averages over the nucleus correlations curves.

### Analytical Model

The fitting was made using the Levenberg-Marquardt algorithm with the nlsLM function from the minpack.lm R package. The confidence intervals of estimated values were computed by 100 bootstrap samples^62^.

### Microscopy

Labeled cells were images by Structured Illumination Microscopy (SIM) on a Zeiss Elyra PS.1 microscope with a 63X oil immersion objective (NA=1.4). Photodetection was made by an EMCCD Andor iXon 885 camera. For each focal plane 15 images were taken, for 5 different phases and 3 different orientations of the modulated illumination pattern. To assess the 3D structure of the cells confocal images were taken every 0.12 µm step on the z-direction. The reconstruction process that gives a final super-resolved images was made with Zen Software. The images were segmented to obtain the 3D position of each individual nuclear pore. Sub-resolution fluorescent beads were used to acquire experimental PSFs that were used for the processing of the SIM images.

### Immunochemistry and labeling

After 3h in culture on the microprinted PDMS coverslips, cells were fixed with 4% PFA at room temperature (RT) for 15 min and then permeabilized with 0.2% Triton X-100 in PBS for 5 min at RT and washed 3 times in PBS. The fixed cells were incubated for 30 min in a blocking solution (0.2% tween, 1% BSA, 1% SVF) and labeled for glycoproteins with wheat germ agglutinin (Alexa Fluor 488 WGA) 1µg/ml for 20 min at RT. The cells were washed 3 times in PBS and labeled for F-actin with SiR-Actin 100 nM in PBS overnight at 4°C. The cells were washed 3 times in PBS and DAPI was added at 0.1µg/ml for 30 min at RT. Coverslips were mounted in Vectashield Antifade Mounting Medium.

### Cell culture

C2C12 myoblasts cells were cultured in DMEM (high glucose, pyruvate, glutamax) medium (FBS 15%) and 1% pen-strep. Cells were plated on the micro-printed PDMS coated coverslips for 3h at 37°C at a density of 5000 cells /cm^2^.

### Microcontact printing

To make the rectangular-shaped micropatterns, we designed rectangular silicon/SU8 mould generated by a photolithographic process. Rectangles are 700-1800µm^2^ with aspect ratio 1:8.

This mould was used to shape a Polydimethylsiloxane stamp (PDMS, 1:10 ratio, Sylgard 184 Dow Corning). After an overnight curing process at 60°C, the PDMS stamp was peeled off the mould and washed for 15 minutes using an ultrasonic bath in a 30% water/70% ethanol solution. The structured PDMS stamp was dried with clean dry air and inked with a fibronectin solution from bovine plasma (FN) at 50 µg/mL for 45 min at room temperature. Meanwhile, a thin layer (10 µm) of PDMS (1:10 ratio) was spin-coated on high performance cover glasses (Carl Zeiss). PDMS coated glass was exposed to UV/O3 irradiation for 7 min. The FN-coated stamp was washed with PBS, dried with clean dry air and then placed in contact with the PDMS coated glass coverslip for a few seconds. The surface was passivated for 45 min with a 0.2% solution of Pluronic F-127. The stamped cover glasses were washed 3 times with PBS and cell culture DMEM was added immediately prior to the experiment.

## Notes

### Competing Interest Statement

The authors have declared no competing interest.

