## Supplementary Material for "Nuclear pore complex distribution on the nuclear envelope: insights into curvature, chromatin, and actin contributions"

**[+] Author contributions:**

These authors contributed equally to this work.

### Supplementary Material

**Curvature measurement**

Note that surfaces are described by two principal curvatures ($\kappa_1$ and $\kappa_2$) representing the maximum and minimum curvatures at a given point on the surface. The surface is well defined by principles curvatures or equivalently, by Gaussian ($K = \kappa_1 \kappa_2$) and average curvatures ($H = \frac{1}{2}(\kappa_1 + \kappa_2)$). Note, we used the mathematical convention for principal curvature signs, *i.e.*, curvature is positive when the normal vector points toward the center of the osculating circle (*e.g.* a sphere has negative curvature). The sign of the Gaussian curvature is not a convention and is meaningful, K>0 indicates that the surface is locally a sphere, K=0 a cylinder (or flat) and K<0 indicates hyperboloid surface (*e.g.* saddle point).

We compared two methods to compute curvatures in combination with two different interpolation methods. We generated a CSR distribution of points at the surface of a hemi-sphere of radius 6.2µm (**see methods**) and reconstructed the surface either with the triangulation-based cubic interpolation or bi-harmonic spline interpolation not based on triangulation (as we did in figure 1-3). Next, we computed curvatures either using the eigenvalues of the Hessian matrix of the surface, or using the Laplace-Beltrami operator on a triangularized version of the surface (**see methods and S10A**). The Hessian matrix requires computing derivatives of the surfaces. Because our surface is discrete, we relied on the Finite Difference Method (FDM) (**see methods**). Our analysis show that cubic interpolation generates triangular edges leading to artifactual curvatures, thus confirming that biharmonic spline interpolation is more accurate than cubic (**S10B**). Additionally, estimation of curvatures by FDM generates significant errors in regions with high slopes while the Laplace-Beltrami operator is robust over a high range of slopes (**S10B**). Furthermore, we compared FDM and Laplace-Beltrami method on simulated perfect hemi-sphere of radius 1µm to drop out the effect of the interpolation. Once again, Laplace-Beltrami demonstrated its robustness compared to FDM (**see methods and table S1**). Laplace-Beltrami method can be affected by the presence of negative weights (see methods), which are the sign of a locally non-Delaunay triangularization, resulting in a poorer meshing and could generate numerical instability[1,2]. More dramatically, we can observe locally $H^2$-K<0 which is mathematically impossible. However, we measured that the proportions of these two artifacts on our experimental surfaces are low (<1% and <0.1% respectively).

| *Pixel size* | Method | 0% | 5% | 10% | 20% | 50% | 80% | 90% | 95% | 100% |
|---|---|---|---|---|---|---|---|---|---|---|
| *100nm* | FDM | 10% | 16% | 19% | 29% | 73% | 173% | 221% | 240% | 242% |
| | LB | <1% | <1% | 2% | 3% | 6% | 9% | 12% | 28% | 141% |
| *50nm* | FDM | 5% | 12% | 18% | 35% | 112% | 337% | 489% | 637% | 798% |
| | LB | <1% | <1% | <1% | 1% | 3% | 4% | 5% | 17% | 864% |
| *40nm* | FDM | 4% | 11% | 18% | 35% | 124% | 386% | 616% | 870% | 1179% |
| | LB | <1% | <1% | <1% | 1% | 2% | 4% | 4% | 15% | 1087% |
| *30nm* | FDM | 3% | 10% | 18% | 36% | 137% | 485% | 834 | 1139 % | 1872% |
| | LB | <1% | <1% | <1% | <1% | 2% | 3% | 3% | 12% | 1309% |

**Table S1**. Distribution of absolute relative errors of FDM and Laplace-Beltrami method on the principal curvature $\kappa_1$ on perfect hemi-sphere of radius 1µm for different pixel sizes. Columns indicate the quantiles of the errors distribution and values are the corresponding errors in %. For our experimental pixel size of 40nm, 90% of the pixels have a relative error inferior to 10% with the Laplace-Beltrami method while with FDM less then 5% of the pixels have a relative inferior to 10%.

### Supplementary Figures

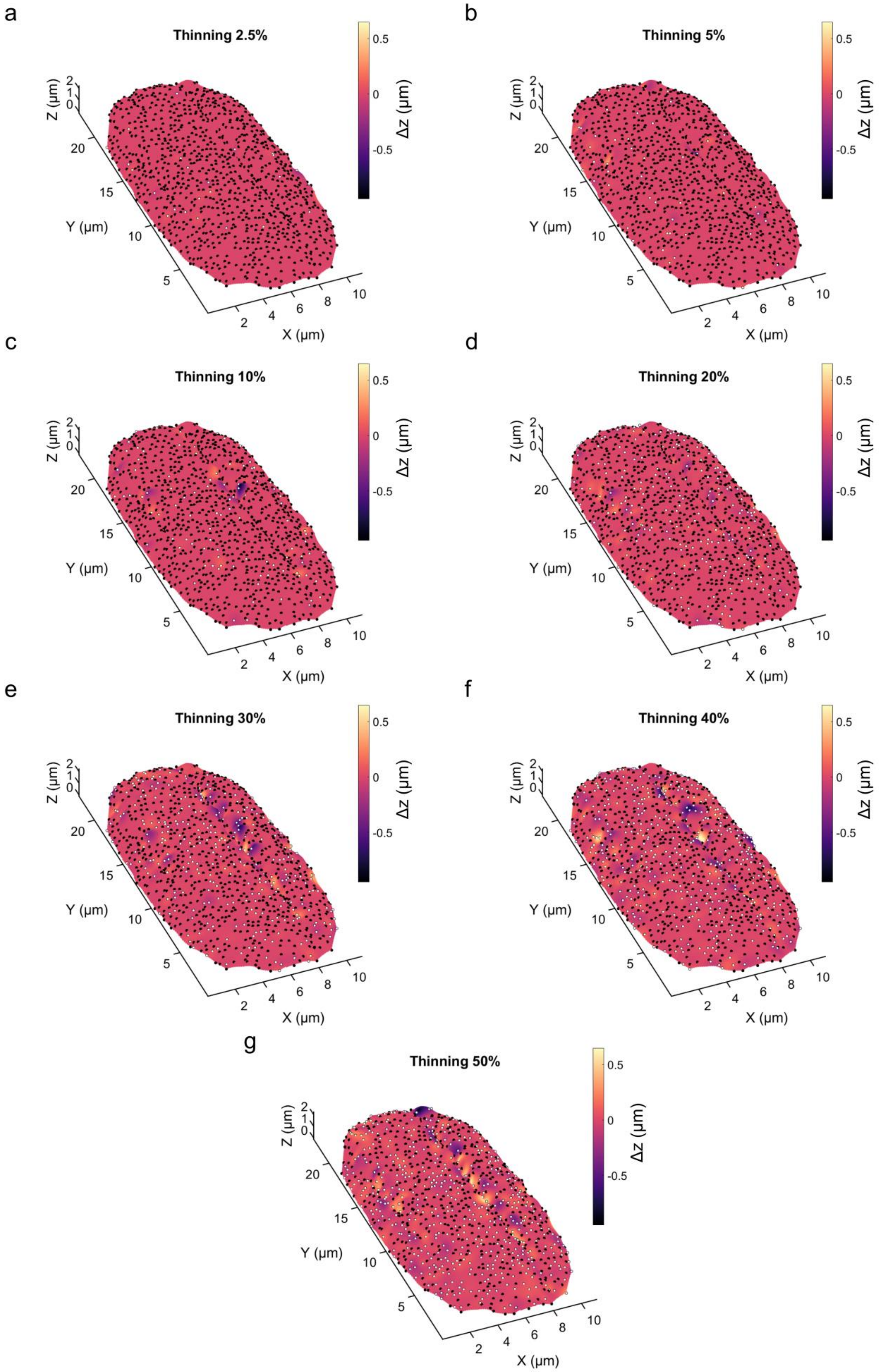
a
Thinning 2.5%
b
Thinning 5%
c
Thinning 10%
d
Thinning 20%
e
Thinning 30%
f
Thinning 40%
g
Thinning 50%
X (µm)
Y (µm)
Z (µm)
Δz (µm)

**Figure S1**. Nuclear surface with color code representing the surface error Δz between reference surface reconstructed using all the NPCs and a surface reconstructed after a thinning of 2.5% (**a**), 5% (**b**), 10% (**c**), 20% (**d**), 30% (**e**), 40% (f) or 50% (**g**) of the points. The display surface is the one reconstructed after thinning. Black dots represent conserved NPCs and white dots represent thinned NPCs.

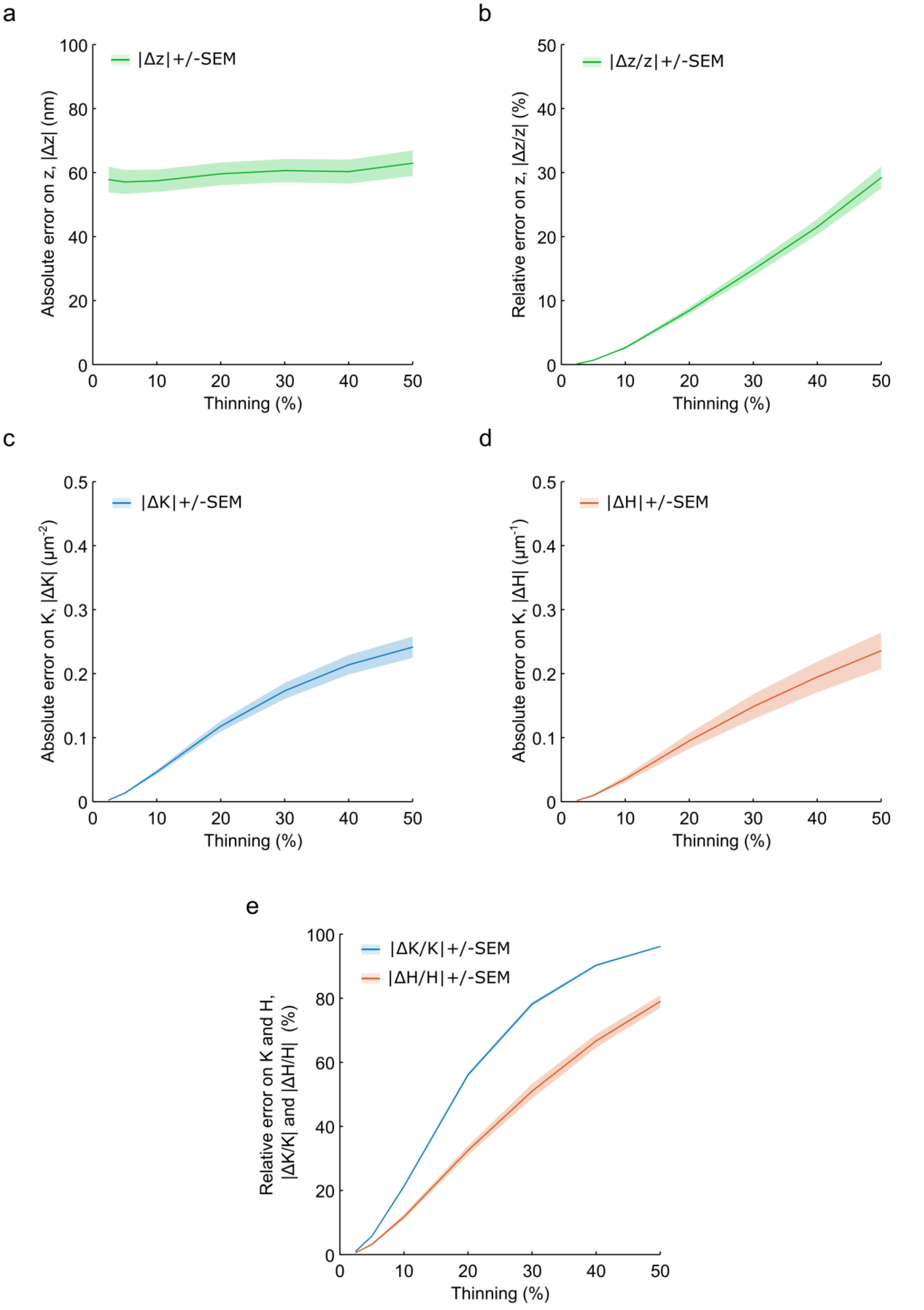


**Figure S2**. **a**. Absolute surface error Δz as a function of the thinning. Absolute surface error is evaluated as the median of |Δz| evaluated over the full nuclear surface. **b**. Relative surface error $\Delta z/z$ as a function of the thinning. Relative surface error is evaluated as the median of $|\Delta z/z|$ evaluated over the full nuclear surface. **c**. Absolute Gaussian curvature error $\Delta K$ as a function of the thinning. Absolute Gaussian curvature error is evaluated as the median of $|\Delta K|$ evaluated over the full nuclear surface. **d**. Absolute average curvature error $\Delta H$ as a function of the thinning. Absolute average curvature error is evaluated as the median of $|\Delta H|$ evaluated over the full nuclear surface. **e**. Relative Gaussian curvature error $\Delta K/K$ (blue) and relative average curvature error $\Delta H/H$ (red) as a function of the thinning. Relative Gaussian and average curvature errors are evaluated as the median of $|\Delta K/K|$ and $|\Delta H/H|$ respectively, evaluated over the full nuclear surface. In a-e, errors are averaged over 57 nuclei ± SEM.

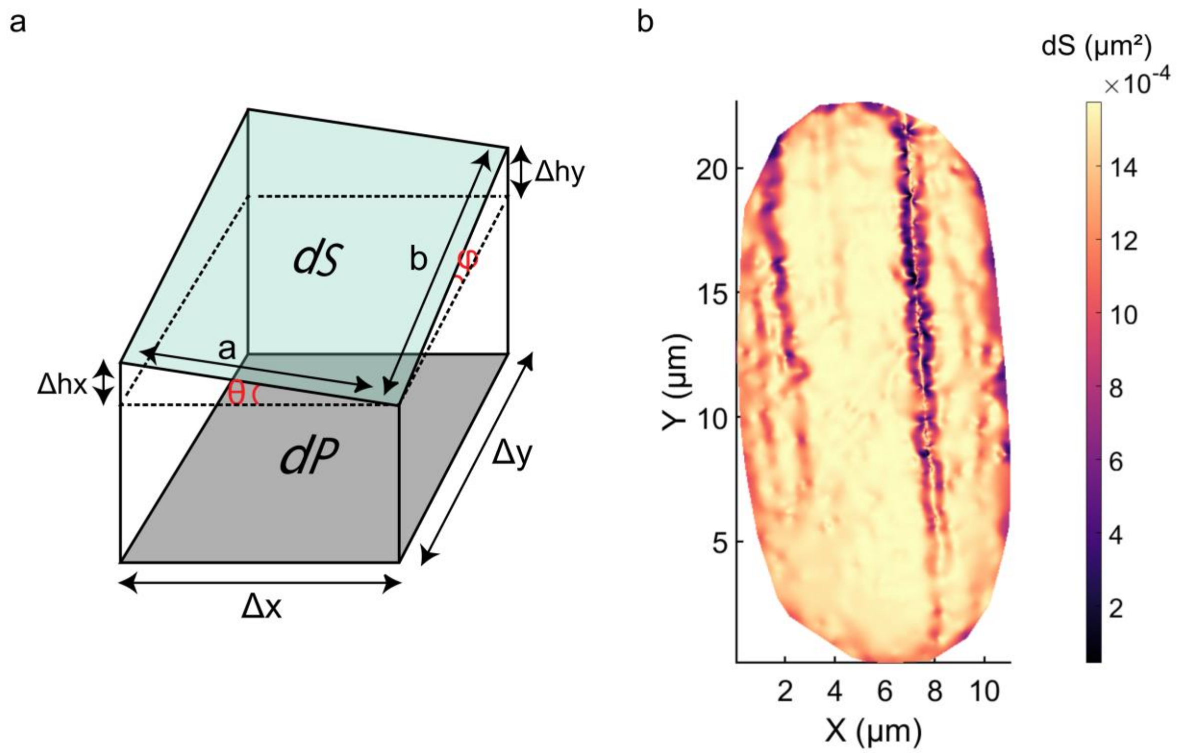


**Figure S3**. **a**. The actual surface element dS can be computed from the projected surface element dP using the gradient of the surface (see methods). **b**. Example of surface element dS map. CSR simulations are achieved using these maps, each surface element has a probability to receive a point directly proportional to its surface dS.

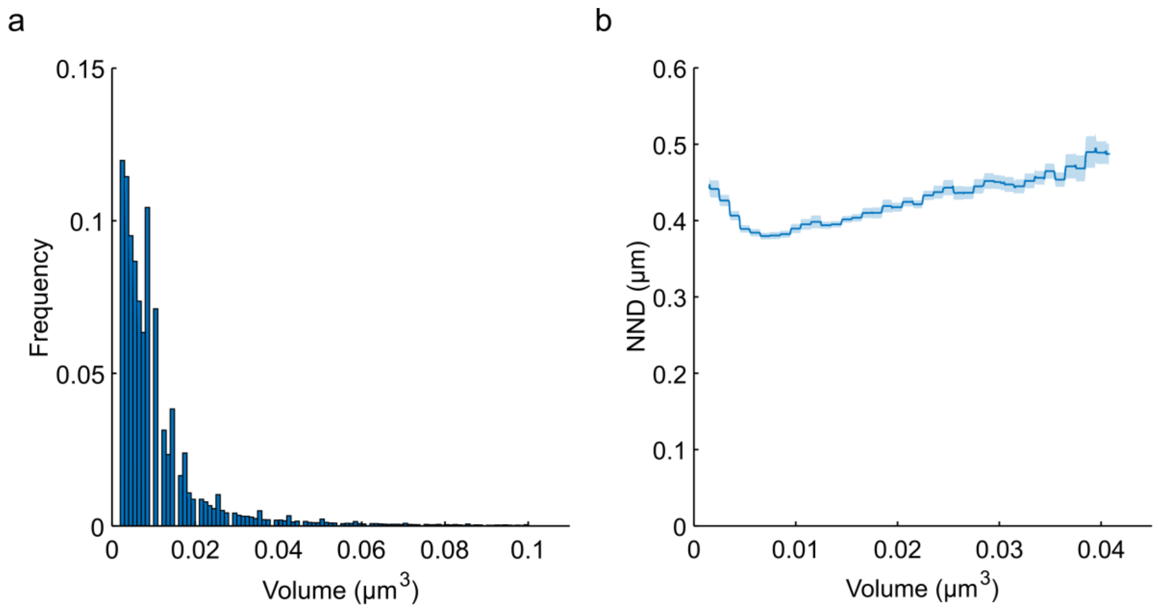


**Figure S4**. **a**. Histogram of segmented NPCs volume from 57 nuclei. **b**. NND as a function of the NPC volume. Solid line is a sliding average ± SEM.

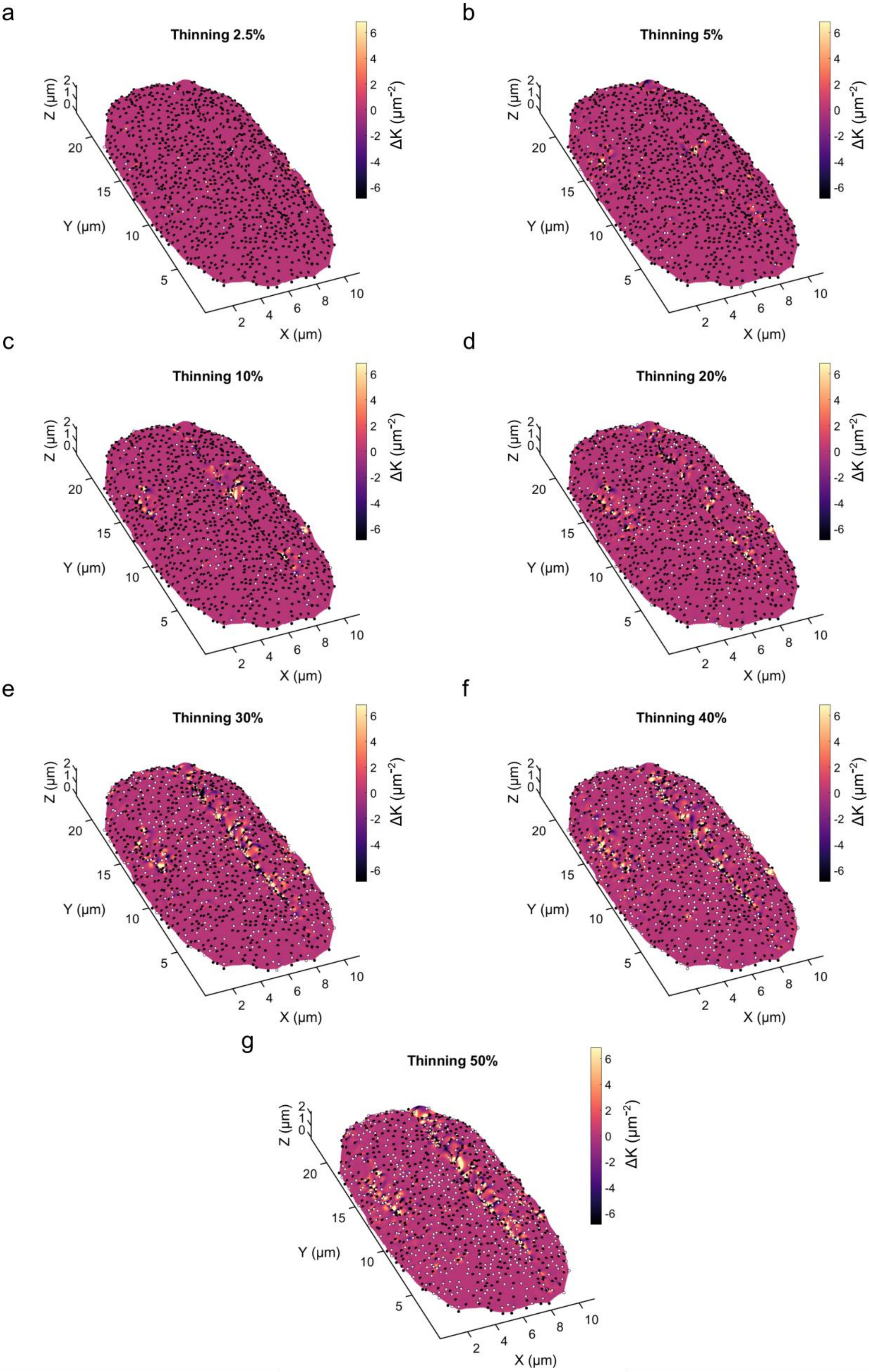
a
Thinning 2.5%
b
Thinning 5%
c
Thinning 10%
d
Thinning 20%
e
Thinning 30%
f
Thinning 40%
g
Thinning 50%
ΔK (μm⁻²)
X (μm)
Y (μm)
Z (μm)

**Figure S5**. Nuclear surface with color code representing the Gaussian curvature error ΔK between reference surface reconstructed using all the NPCs and a surface reconstructed after a thinning of 2.5% (**a**), 5% (**b**), 10% (**c**), 20% (**d**), 30% (**e**), 40% (f) or 50% (**g**) of the points. The display surface is the one reconstructed after thinning. Black dots represent conserved NPCs and white dots represent thinned NPCs.

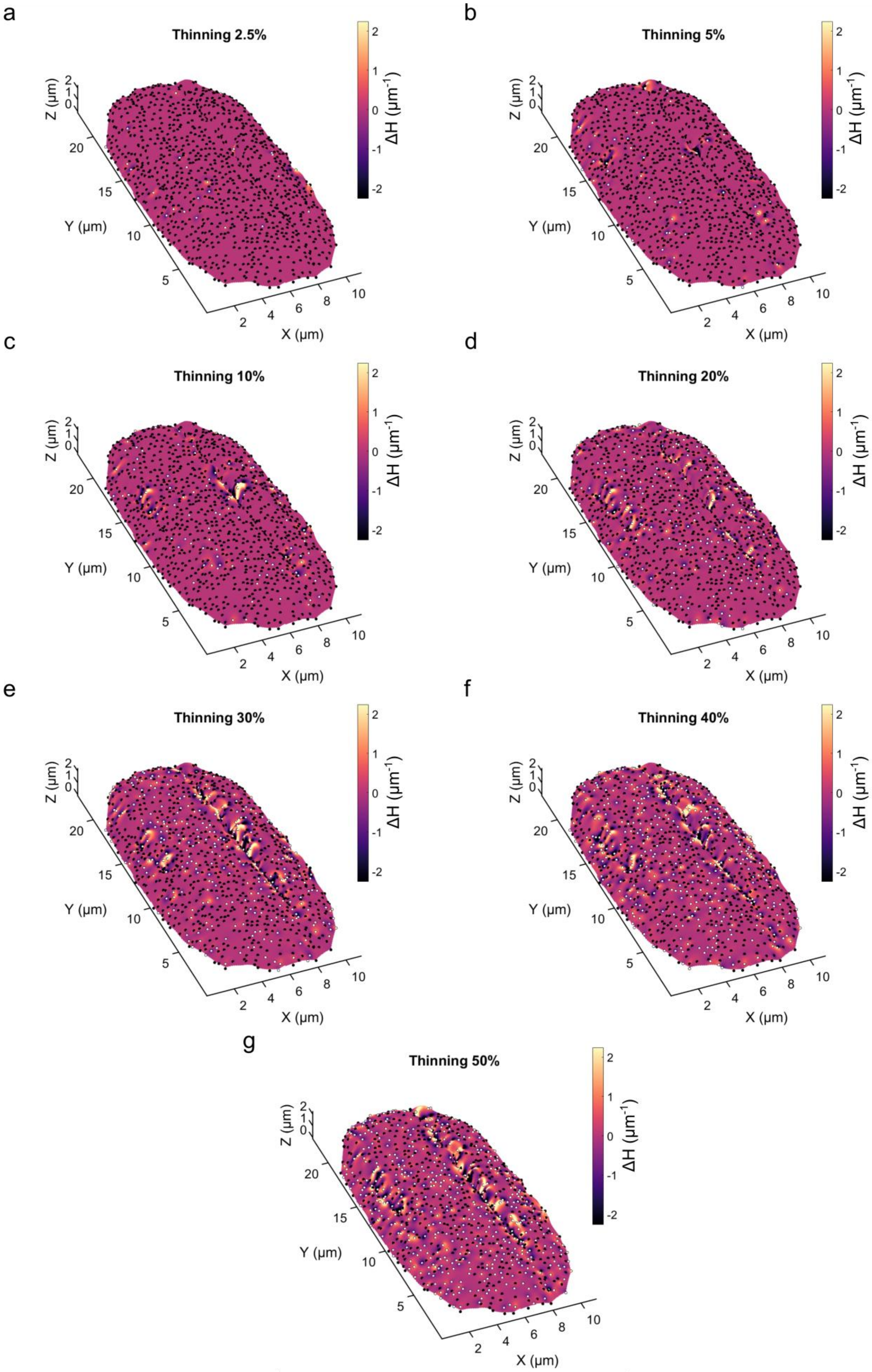

a
Thinning 2.5%
b
Thinning 5%
c
Thinning 10%
d
Thinning 20%
e
Thinning 30%
f
Thinning 40%
g
Thinning 50%
Z (µm)
Y (µm)
X (µm)
ΔH (µm⁻¹)

**Figure S6**. Nuclear surface with color code representing the mean curvature error ΔH between reference surface reconstructed using all the NPCs and a surface reconstructed after a thinning of 2.5% (**a**), 5% (**b**), 10% (**c**), 20% (**d**), 30% (**e**), 40% (f) or 50% (**g**) of the points. The display surface is the one reconstructed after thinning. Black dots represent conserved NPCs and white dots represent thinned NPCs.

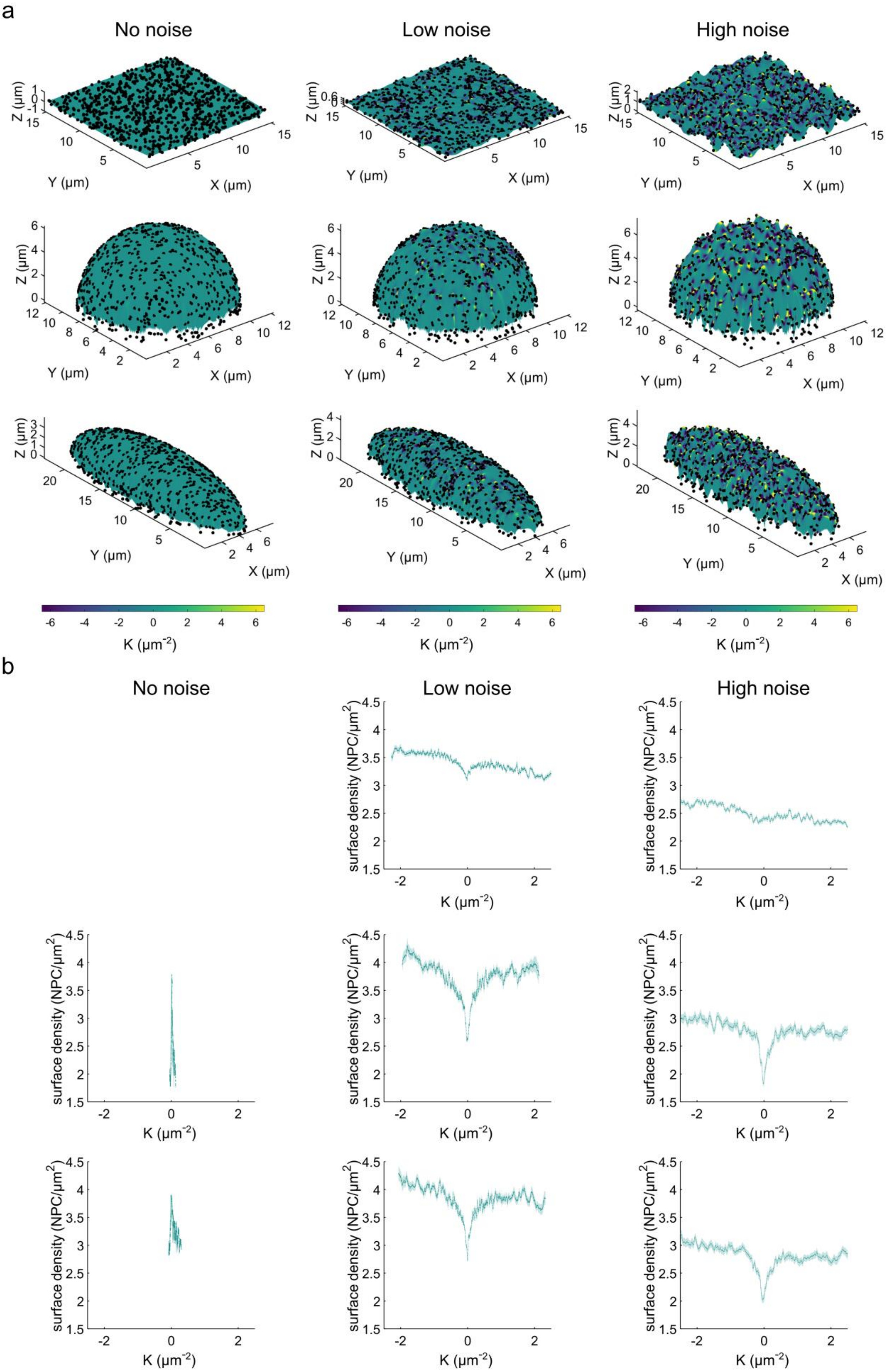

a
No noise
Low noise
High noise
Z (μm)
Y (μm)
X (μm)
K (μm$^{-2}$)
b
No noise
Low noise
High noise
surface density (NPC/μm$^2$)
K (μm$^{-2}$)

**Figure S7**. **a**. Surface reconstruction by interpolation of simulated NPCs (Matérn II hard core simulation) on different geometries (flat square, hemi-sphere or hemi-ellipsoid) for different levels of noise on the z coordinates (no noise, low noise or high noise). Black dots represent simulated NPCs and color code represents the Gaussian curvature K. Simulations have NPC density similar to the one experimentally measured. **b**. NPC surface density as a function of the Gaussian curvature K. Solid line represents an average over 100 simulations and the shade the SEM. Moreover, and similarly to experimental data, a 16x16 (640x640µm) binning is applied.

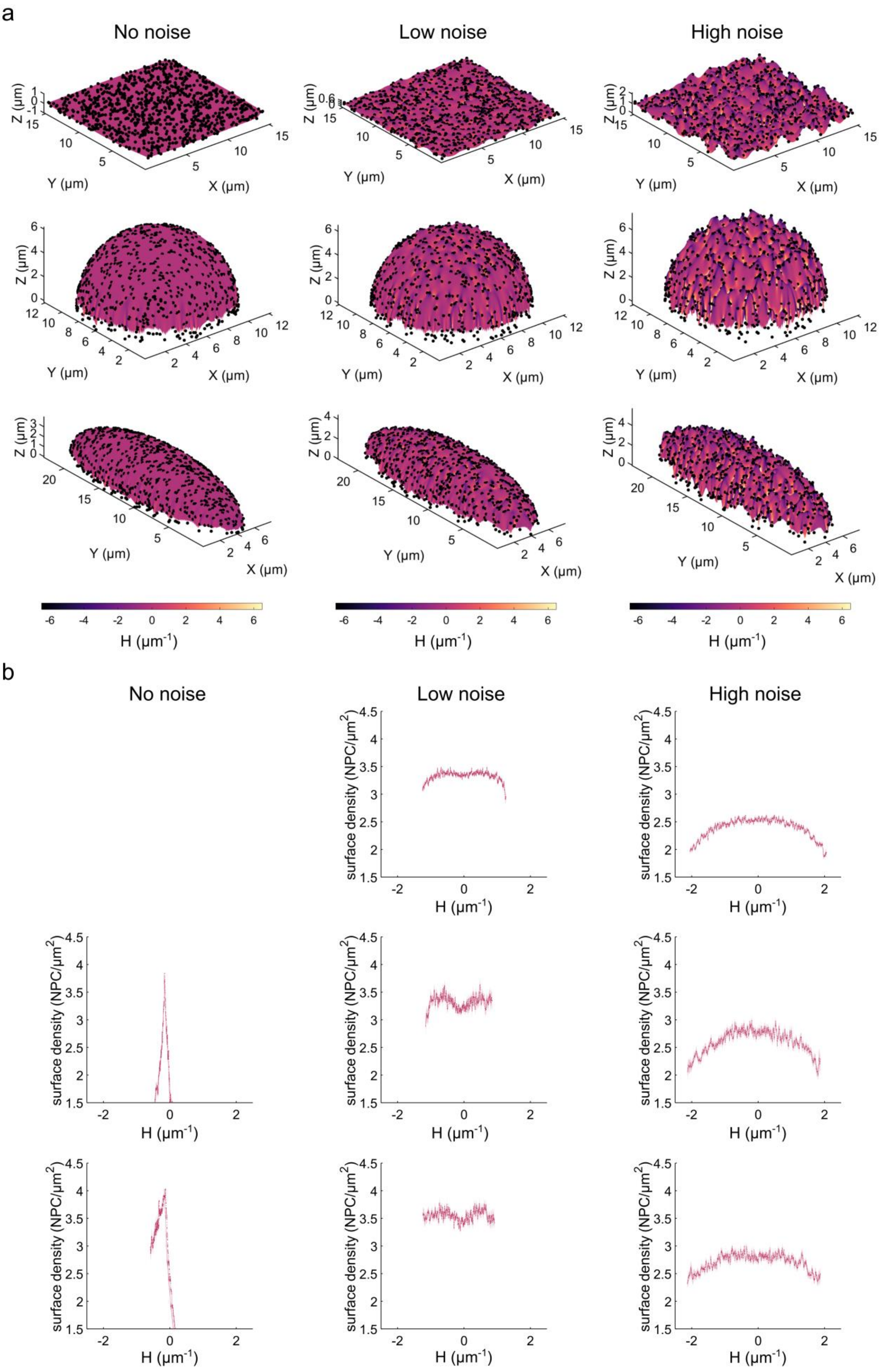
a
No noise
Low noise
High noise
Z (μm)
Y (μm)
X (μm)
H (μm$^{-1}$)
b
No noise
Low noise
High noise
surface density (NPC/μm$^2$)
H (μm$^{-1}$)

**Figure S8. a**. Surface reconstruction by interpolation of simulated NPCs (Matérn II hard core simulation) on different geometries (flat square, hemi-sphere or hemi-ellipsoid) for different levels of noise on the z coordinates (no noise, low noise or high noise). Black dots represent simulated NPCs and color code represents the mean curvature H. Simulations have NPC density similar to the one experimentally measured. **b**. NPC surface density as a function of the mean curvature H. Solid line represents an average over 100 simulations and the shade the SEM. Moreover, and similarly to experimental data, a 16x16 (640x640µm) binning is applied.

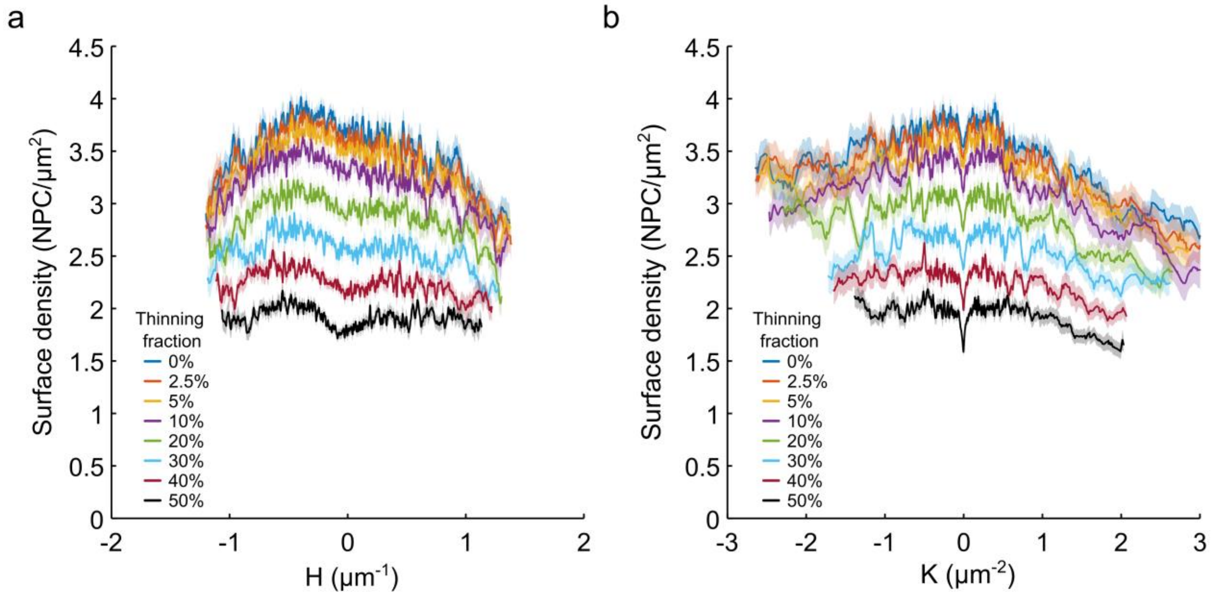


**Figure S9. a**. Experimental NPC surface density as a function of the mean curvature H for different thinning fractions. **b**. Experimental NPC surface density as a function of the Gaussian curvature K for different thinning fractions. In a-b, Solid lines represent an average over 57 nuclei and the shade the SEM. The thinning reduces the total number of points leading to the translation of the curves along the y-axis. As in figure 4, a 16x16 (640x640μm) binning is applied.

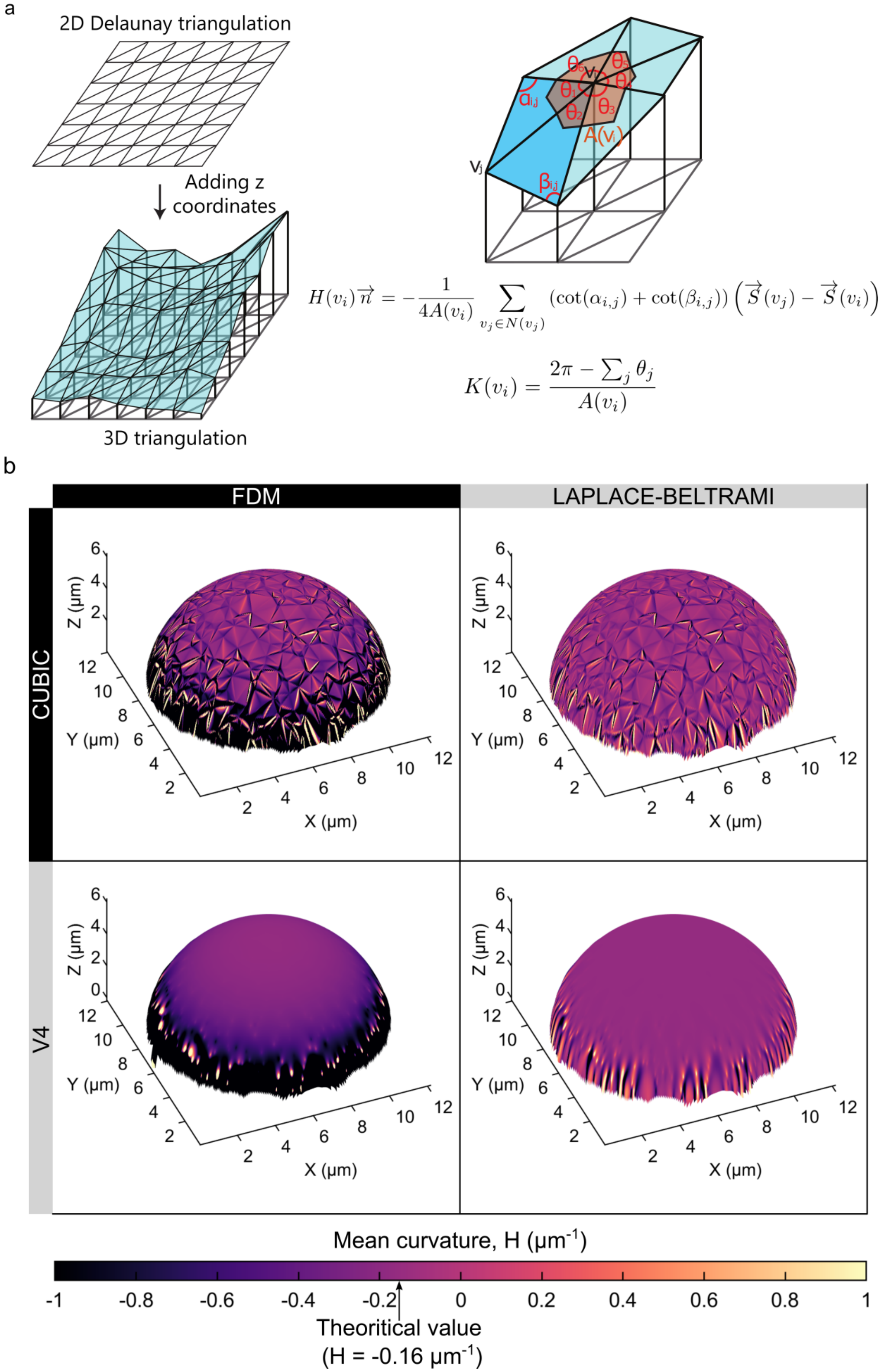
a
2D Delaunay triangulation
Adding z coordinates
3D triangulation
$\alpha_{i,j}$
$\beta_{i,j}$
$v_i$
$v_j$
$\theta_1$ $\theta_2$ $\theta_3$ $\theta_4$ $\theta_5$ $\theta_6$
$A(v_i)$
$$H(v_i)\vec{n} = -\frac{1}{4A(v_i)} \sum_{v_j \in N(v_j)} (\cot(\alpha_{i,j}) + \cot(\beta_{i,j})) \left(\vec{S}(v_j) - \vec{S}(v_i)\right)$$
$$K(v_i) = \frac{2\pi - \sum_j \theta_j}{A(v_i)}$$
b
FDM
LAPLACE-BELTRAMI
CUBIC
V4
Z (µm)
Y (µm)
X (µm)
Mean curvature, H (µm$^{-1}$)
-1 -0.8 -0.6 -0.4 -0.2 0 0.2 0.4 0.6 0.8 1
Theoritical value
(H = -0.16 µm$^{-1}$)

**Figure S10**. **a**. Illustration of the scheme to compute Gaussian and mean curvatures. Starting from 2D trivial Delaunay triangulation, we added the z coordinates to re-create the surface keeping the triangulation. From this 3D triangulated surface, we computed curvatures using angles and surfaces illustrated on the figure (see methods). **b**. Comparison of surface interpolation methods (cubic or biharmonic splines (V4)) in combination with different methods to compute Gaussian curvatures (FDM or Laplace-Beltrami-operator) from points generated at the surface of a hemi-sphere of radius r=6.20µm. The color code indicates the theoretical mean curvature equal to $H=-0.16\mu m^{-1}$.
